# An area postrema GLP-1R to NTS pathway dissociates semaglutide-induced satiation, nausea and reduced food motivation from satiety

**DOI:** 10.64898/2026.08.10.744052

**Authors:** Lauren A. Jones, Ella Cross, Yuqi Song, Nicholas Monaco, Phoebe Claxton, Yuan Yu, Stefan Trapp, Alice Adriaenssens, Daniel I. Brierley

**Affiliations:** UCL Centre for Cardiovascular and Metabolic Neuroscience, Department of Neuroscience, Physiology and Pharmacology, University College London, Gower St, WC1E 6BT, United Kingdom

## Abstract

The GLP-1-based anti-obesity drug semaglutide lowers bodyweight primarily by increasing satiation and satiety, whilst also reducing motivation to consume high calorie foods, and commonly causing nausea. The brainstem dorsal vagal complex (DVC) has been identified as a key mediator of these phenotypic components of semaglutide’s anorectic effect. However, which DVC neuron populations are recruited to mediate these components and whether they are dissociable, are translationally important but unresolved questions. We addressed these in mice using metabolic and behavioural phenotyping to resolve the distinct anorectic components elicited by the acute administration of semaglutide. We combined this phenotyping approach with activity-dependent genetic labelling to identify semaglutide-recruited neuron populations (‘Sema-TRAP’) and chemogenetic manipulation to interrogate their contribution to these effects.

Semaglutide potentiated satiation and satiety, caused behavioural proxies of nausea, and suppressed motivation to consume high-fat, high-sugar Western diet, in a largely sex-independent manner. It activated a substantial proportion of GLP-1R-expressing neurons in the brainstem area postrema (AP), but surprisingly most semaglutide-activated neurons in the nucleus tractus solitarius (NTS) did not express GLP-1R. Chemogenetic reactivation of Sema-TRAP neurons in the NTS alone was sufficient to recapitulate the effects of semaglutide on satiation, nausea, motivation for Western diet, and bodyweight. Knockdown of GLP-1R expression in the AP prior to Sema-TRAPing prevented the recruitment of Sema-TRAP^NTS^ neurons which elicited these effects on satiation, nausea and motivation. AP GLP-1R knockdown also abolished semaglutide’s effect on satiation and motivation, while effects on satiety and bodyweight were preserved. These data demonstrate that semaglutide suppresses eating via distinct behavioural mechanisms, at least some of which are mediated by dissociable circuits. GLP-1R populations outside the DVC, or distinct subpopulations within the non-GLP-1R NTS neurons recruited downstream of GLP-1R^AP^, may therefore represent therapeutic targets to tune GLP-1-based obesity drugs towards a better-tolerated effect profile.

## Introduction

Glucagon-like peptide-1 receptor agonists (GLP-1RAs), such as semaglutide, are highly effective treatments for diabetes and obesity, due to their robust beneficial effects on glucose homeostasis and bodyweight^1^. However, nausea and other adverse effects are common and substantially limit adherence and effectiveness^2^. Beyond their well-established actions to promote satiation and satiety, GLP-1RAs are reported to selectively reduce preference for highly palatable foods, and attenuate motivation to consume both food and non-food rewards, including alcohol and recreational drugs^3,4^. Despite these diverse behavioural effects, the precise neural circuitry through which GLP-1RAs modulate satiation, satiety, nausea and reward-related behaviours remain incompletely understood. Identifying the neuronal populations mediating these behavioural effects, and whether the beneficial phenotypic components are mediated by populations distinct from those which mediate compliance-limiting side effects, could offer opportunities to tune next-generation GLP-1-based obesity drugs towards a better-tolerated effect profile.

Studies using *Glp1r*-Cre mice have identified the dorsal vagal complex (DVC), comprising the area postrema (AP), nucleus tractus solitarius (NTS) and dorsal motor nucleus of the vagus, as a key locus for the anorectic and aversive effects of semaglutide^5–7^. While activation of either AP or NTS GLP-1R neurons is sufficient to reduce food intake and body weight^5^, their individual necessity for the therapeutic and/or undesirable effects of semaglutide is contentious. Accumulating evidence suggests that semaglutide and other GLP-1RAs may only recruit subsets of AP and NTS GLP-1R neurons^8–10^, in addition to unidentified non-GLP-1R populations, highlighting the limitations of receptor-defined cell type-specific approaches for identifying the neuronal ensembles engaged by semaglutide *in vivo*. Accordingly, despite several region-specific GLP-1R knockdown, inhibition and neuronal ablation studies^5–7^, the neuronal populations required for the specific anorectic effects of semaglutide remain incompletely defined. In particular, whether NTS GLP-1R neurons are recruited directly by semaglutide or engaged downstream of other GLP-1R populations, and whether distinct DVC neuronal circuits mediate its homeostatic, hedonic and/or aversive effects, remains unresolved. A major hurdle in this respect is the marked heterogeneity of cell-types within the DVC, with at least 10 largely distinct populations of anorectic neurons identified in the NTS alone, and GLP-1R expression identified in multiple populations^11–14^.

Beyond its effects on homeostatic control of food intake, semaglutide has been shown to reduce preference for high-fat, high-sugar foods (sometimes termed the ‘Western’ diet), and attenuate reward-motivated behaviours in people with substance use disorders^3,4,15^. Consistent with this, recent preclinical studies have demonstrated that pharmacological, chemogenetic and/or optogenetic manipulation of GLP-1R neurons in mesolimbic regions including the ventral tegmental area (VTA)^16^, lateral septum (LS)^17^, central amygdala (CeA)^3,18^ is sufficient to suppress reward-motivated behaviours. However, it is unclear whether any of these are direct sites of action for semaglutide specifically, or if these circuits are recruited indirectly by accessible upstream populations of GLP-1R neurons.

Cell type-specific approaches to study GLP-1R-expressing neuronal populations have yielded many valuable insights into central GLP-1 system signalling in the context of eating behaviour and metabolic control^5,18,19^. However, the extent to which these receptor populations and/or circuits are accessible to and recruited by peripherally-administered semaglutide is unclear. Similarly, it remains unclear which central GLP-1R populations mediate the homeostatic, hedonic and/or aversive effects of semaglutide and to what extent they are dissociable.

To address these unknowns, we developed an activity-dependent genetic tagging approach to anatomically and functionally characterise semaglutide-recruited neurons in the DVC, agnostic to cell-type. We used this model to comprehensively interrogate the role of semaglutide-recruited DVC neurons in the homeostatic, hedonic and aversive phenotypic components of semaglutide’s effects on eating behaviours.

## Results

### Semaglutide potentiates satiation, satiety, and nausea, in a largely sex-independent manner

The effects of semaglutide on food intake and bodyweight have been characterised in several model species, but comparative assessments of its distinct effects on eating behaviours and whole-body metabolism remain limited. We sought to test whether components of the semaglutide-induced behavioural and metabolic phenotype can be dissociated at the level of specific neuronal circuits. To enable this, we first needed to comprehensively characterise the behavioural and metabolic phenotype of semaglutide in mouse, to serve as a reference dataset against which the circuit basis of each discrete phenotypic component could be dissected.

We used a moderate 60 µg/kg (∼15 nmol/kg) acute dose of semaglutide, which suppresses eating behaviours and reduces bodyweight to a translationally relevant extent, and enables comparisons to previous literature^5,10,20,21^. Given emerging evidence for quantitative sex-differences in the effects of some GLP-1RAs in humans and pre-clinical models, but a general paucity of pre-clinical data from female animal models in this context^22^, we conducted all our experiments in sex-balanced cohorts.

Administration of semaglutide 30 minutes prior to dark onset potently reduced chow intake during hours 2-6. There was no compensatory increase in food consumption after this period, with a 54% reduction in cumulative intake over the full circadian cycle (Fig. 1a-c, EDFig. 1f-g). Energy expenditure was also reduced over hour 2-6 (Fig. 1d, EDFig. 1a), but despite this reduction, mice were in overall negative energy balance during this period (Fig. 1e). The reduction in energy expenditure persisted for the entire dark phase (EDFig. 1b-d). The cumulative energy deficit accounted for the observed loss in bodyweight (Fig 1f). Concurrent with the anorectic effect of semaglutide, we saw a reduction in RER during hours 3-5, consistent with a shift towards lipid oxidation (EDFig. 1e). There was a reduction in locomotor activity throughout most of the dark phase (Fig. 1g), and mice also consumed very little water until the light phase (Fig. 1h). Analysis of meal patterns during the acute anorectic phase revealed that semaglutide reduced chow intake by potentiating both satiation, evidenced by smaller and shorter meals (Fig. 1i-j) and satiety, evidenced by less frequent meals, longer inter-meal intervals, and longer post-meal durations per calorie consumed (Fig. 1k-l, EDFig. 1h). The majority of semaglutide’s effects on whole body metabolism were sex-independent, with chow intake reduced to a slightly greater extent in male mice, but both sexes exhibited persistent anorectic effects in dark and light phases (EDFig. 1i-k,s-u). Notably, energy expenditure during the acute anorectic phase was only reduced in male mice, but both sexes were still in negative energy balance during this period, which was sufficient to drive weight loss, albeit at a lower level in females (EDFig. 1l-n,v-x). The reduced chow intake during this period was driven by effects on satiation and satiety in both sexes, although meal duration and inter-meal interval were only significantly altered in male mice (EDFig. 1o-r,y-bb).

**Figure 1:**
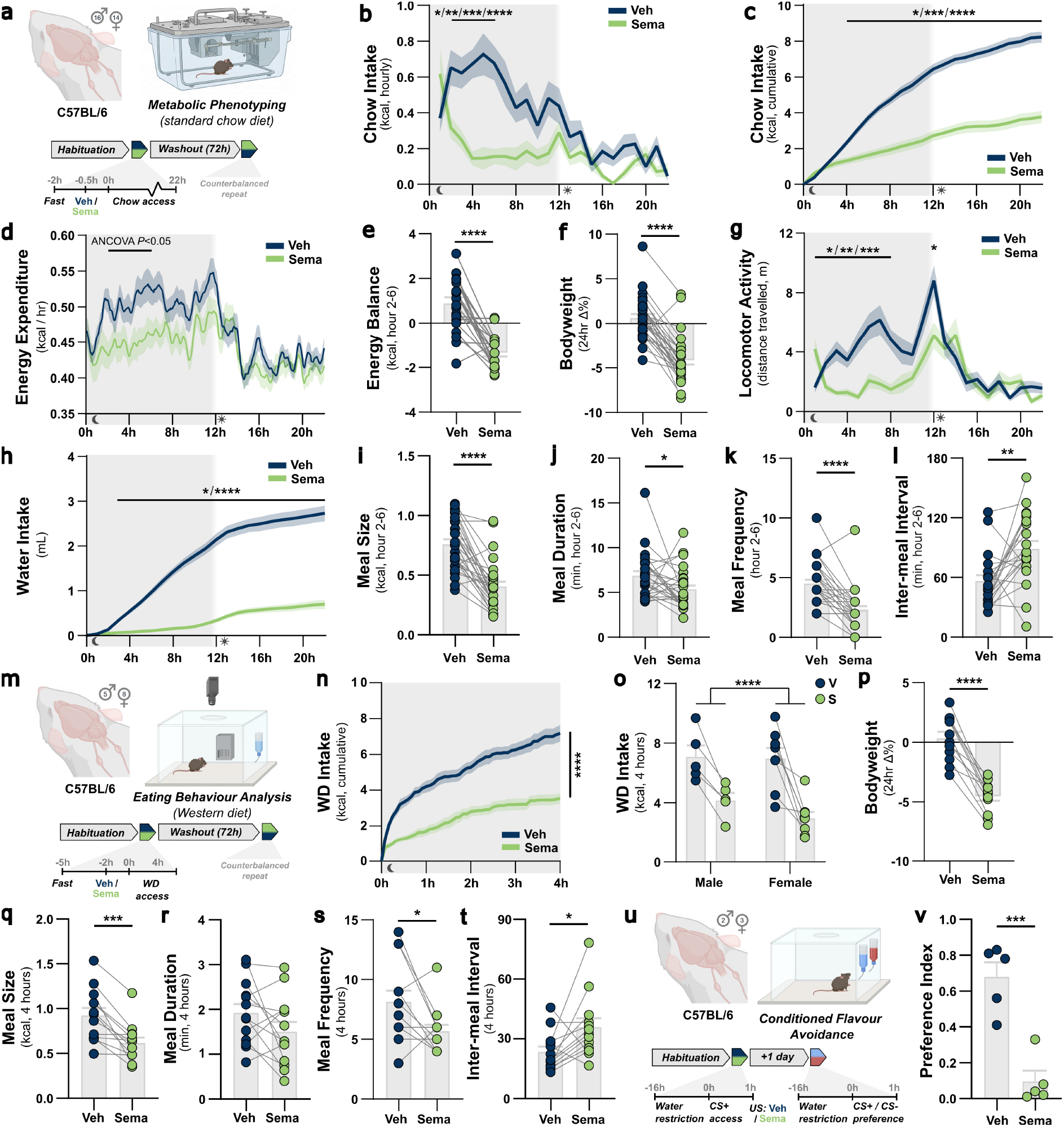
Semaglutide potentiates satiation, satiety, and nausea in a largely sex-independent manner. **a**, experimental model and design for metabolic phenotyping study in chow-fed C57BL/6 mice. **b**, food intake (hourly non-cumulative): drug x hour *F*_(9.9, 285.8)_=4.48, *P*<0.0001. **c,** food intake (hourly cumulative): drug x hour *F*_(22, 638)_=74.1, *P*<0.0001. **d**, energy expenditure plotted as 5 minute bins with 30 minute rolling average (for visualisation purposes only, unsmoothed hourly data used for ANCOVA analysis as fully reported in EDFig. 1a-c). **e**, energy balance (hour 2-6): *W*_(20)_=-219.0, *P*<0.0001. **f**, bodyweight: *t*_(28)_=9.6, *P*<0.0001. **g**, locomotor activity: drug x hour *F*_(10.2, 357.4)_=6.0, *P*<0.0001. **h**, water intake (hourly cumulative): drug x hour *F*_(1.8, 56.4)_=150.1, *P*<0.0001. **i**, meal size (hour 2-6): *t*_(26)_=7.2, *P*<0.0001. **j**, meal duration (hour 2-6): *t*_(26)_=2.3, *P*=0.0328. **k**, meal frequency (hour 2-6): *t*_(29)_=6.2, *P*<0.0001. **l**, inter-meal interval (hour 2-6): *t*_(20)_=3.2, *P*=0.0043. **m**, experimental model and design for high-resolution eating behaviour study in intermittent Western diet (WD)-fed mice. **n**, 4 hour cumulative WD intake (plotted in 5 minute time bins): *t*_(12)_=7.0, *P*<0.0001. **o**, WD intake by sex: drug *F*_(1,11)_=35.8, *P*<0.0001; drug x sex *F*_(1,11)_=0.9, *P*=0.3739. **p**, bodyweight: *t*_(11)_=9.2, *P*<0.0001. **q**, meal size (4 hours): *t*_(12)_=5.0, *P*=0.0003. **r**, meal duration (4 hours): *t*_(12)_=1.8, *P*=0.1048. **s**, meal frequency (4 hours): *t*_(12)_=2.6, *P*=0.0230. **t**, inter-meal interval (4 hours): *t*_(12)_=2.4, *P*=0.0321. **u**, experimental model and design for conditioned flavour avoidance assay. **v**, CFA preference index: *U*=0, *P*=0.0079. All data presented as mean ± sem. * *P*<0.05, ** *P*<0.01, *** *P*<0.001, **** *P*<0.0001.

**Extended Data Figure 1:**
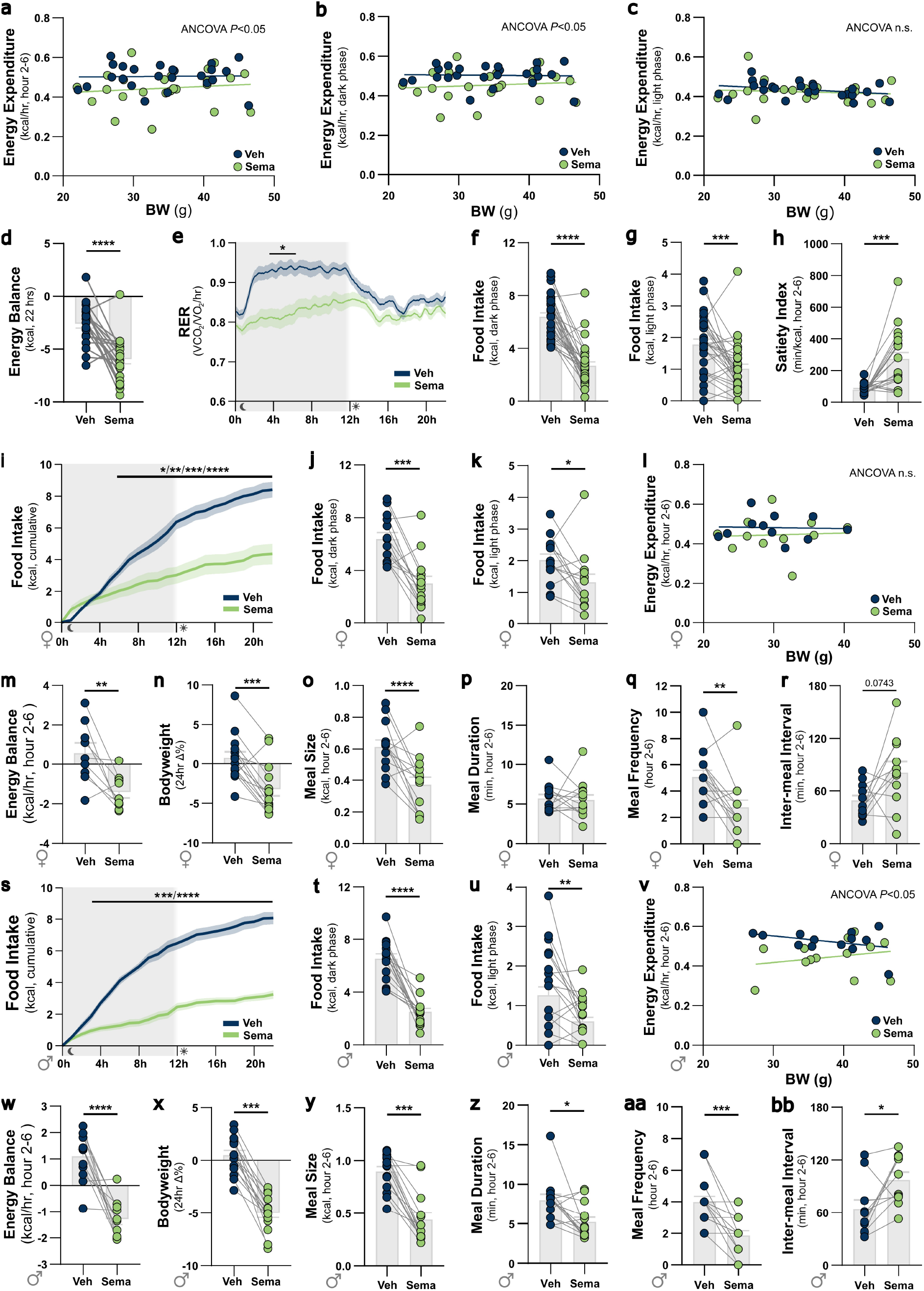
Semaglutide potentiates satiation, satiety, and nausea in a predominantly sex-independent manner. **a**, energy expenditure (hour 2-6) ANCOVA: drug *F*_(1, 45)_=6.8, *P*=0.012. **b**, energy expenditure (dark phase) ANCOVA: drug *F*_(1, 45)_=6.9, *P*=0.012; **c**, energy expenditure (light phase) ANCOVA: drug *F*_(1, 45)_=0.64 *P*=0.429; **d**, energy balance (22 hours): *t*_(21)_=5.4, *P*<0.0001. **e**, RER plotted as 5 minute bins with 30 minute rolling average (for visualisation purposes only, unsmoothed hourly data used for ANOVA analysis): drug x hour *F*_(2.9, 69.2)_=4.8, *P*=0.0045. **f**, food intake (dark phase): *W*_(30)_=-451.0, *P*<0.0001. **g**, food intake (light phase): *t*_(29)_=4.0, *P*=0.0004. **h**, satiety index (hour 2-6): *t*_(20)_=4.7, *P*=0.0002. **i**, food intake (hourly cumulative, females): drug x hour *F*_(2.2, 28.2)_=31.2, *P*<0.0001. **j**, food intake (dark phase, females): *t*_(13)_=5.4, *P*=0.0001. **k**, food intake (light phase, females): *t*_(13)_=2.6, *P*=0.0235. **l**, energy expenditure (hour 2-6, females) ANCOVA: drug *F*_(1, 19)_=1.3, *P*=0.272. **m**, energy balance (hour 2-6, females): *t*_(8)_=3.4, *P*=0.0089. **n**, bodyweight (hour 2-6, females): *t*_(13)_=4.9, *P*=0.0003. **o**, meal size (hour 2-6, females): *t*_(12)_=3.8, *P*=0.0024. **p**, meal duration (hour 2-6; females): *t*_(13)_=0.2, *P*=0.8300. **q**, meal frequency (hour 2-6, females): *t*_(13)_=3.7, *P*=0.0029. **r**, inter-meal interval (hour 2-6, females): *t*_(10)_=2.0, *P*=0.0743. **s**, food intake (hourly cumulative, males): drug x hour *F*_(2.0, 30.0)_=44.2, *P*<0.0001. **t**, food intake (dark phase, males): *t*_(15)_=8.1, *P*<0.0001. **u**, food intake (light phase, males): *t*_(15)_=3.0, *P*<0.0085. **v**, energy expenditure (hour 2-6, males) ANCOVA: drug *F*_(1, 23)_=6.6, *P*=0.0170. **w**, energy balance (hour 2-6, males): *t*_(11)_=3.0, *P*<0.0001. **x**, bodyweight (hour 2-6, males): *t*_(14)_=10.4, *P*<0.0001. **y**, meal size (hour 2-6, males): *W*_(13)_=-91.0, *P*=0.0002. **z**, meal duration (hour 2-6; males): *t*_(14)_=2.9, *P*=0.0117. **aa**, meal frequency (hour 2-6, males): *t*_(15)_=5.2, *P*=0.0001. **bb**, inter-meal interval (hour 2-6, males): *t*_(9)_=2.6, *P*=0.0291. All data presented as mean ± sem. # *P*<0.1, * *P*<0.05, ** *P*<0.01, *** *P*<0.001, **** *P*<0.0001.

Following this comprehensive metabolic phenotyping study of normal chow-fed mice, we next conducted high-resolution eating behaviour analyses to characterise the effects of semaglutide on consumption of a high-fat, high-sugar Western diet (WD). Metabolic phenotyping systems can be susceptible to inaccurate recording of eating bouts, particularly for high-fat diet pellets. This can confound attempts to perform accurate meal pattern analysis in response to acute drug administration or transient circuit manipulations. To overcome this limitation, we developed an acute eating behaviour analysis platform where standard operant chambers were modified to resemble home-cage environments. They were equipped with a highly sensitive food intake monitoring system^23^, and infrared cameras for concurrent video monitoring, enabling accurate, high-resolution assessment of WD meal microstructure during the dark phase, and additional automated behavioural analyses (EDFig. 2a-c).

**Figure 2:**
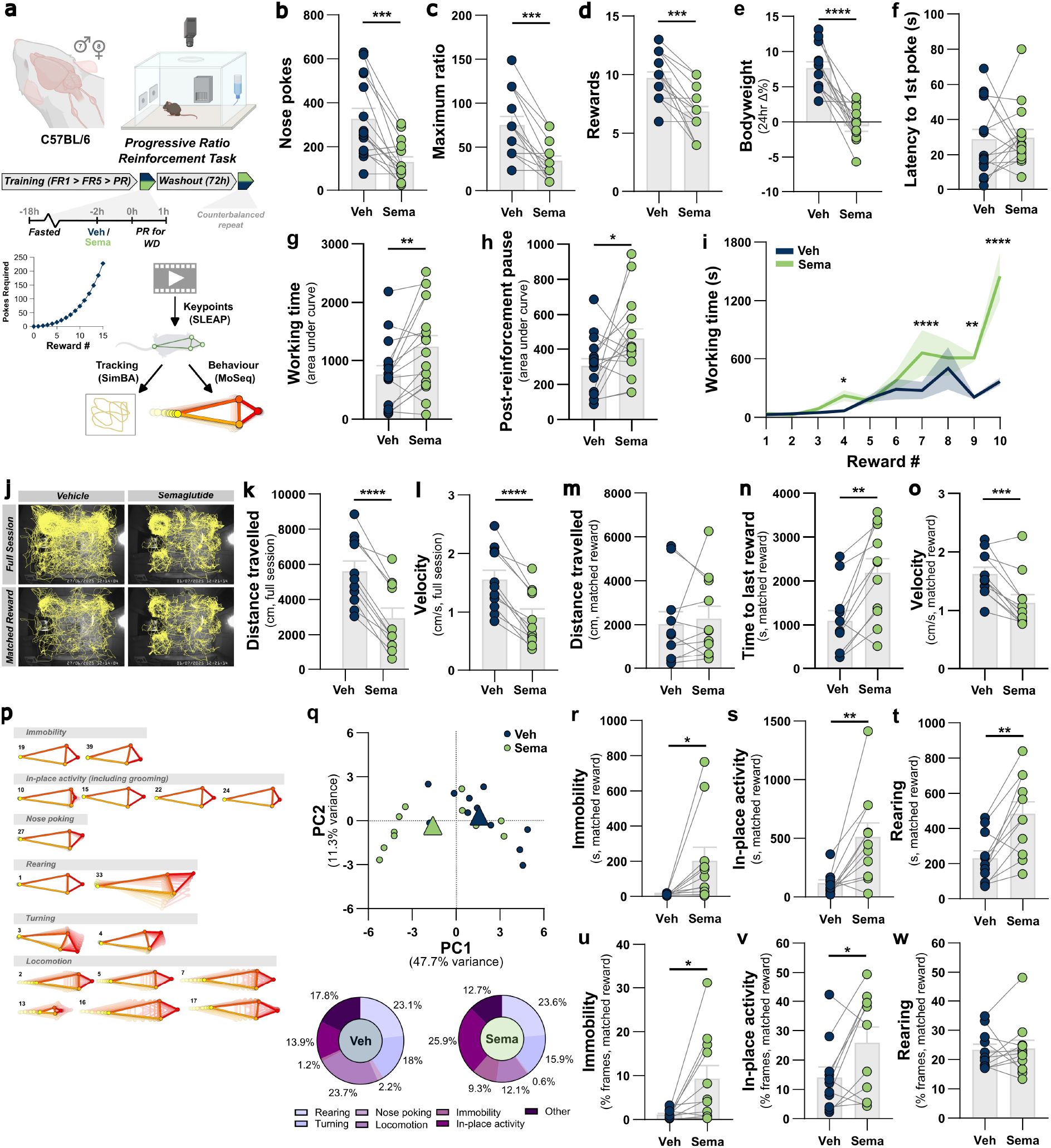
Semaglutide suppresses motivation to consume high-fat high-sugar Western diet. **a**, experimental model and design for progressive ratio (PR) schedule of reinforcement task for WD rewards in C57BL/6 mice, including exponential PR schedule of nose pokes required for to gain successive access to the WD hopper for 7s (1, 3, 6, 10, 16, 23, 32, 43, 57, 74, 94, 119, 149, 185, 228), and software packages used to train video tracking model and analyse behaviour during PR task. **b**, total nose pokes: *t*_(14)_=4.8, *P*=0.0003. **c**, maximum ratio completed: *t*_(14)_=5.3, *P*=0.0001. **d**, total rewards earned: *t*_(14)_=5.3, *P*=0.0001. **e**, 24hr bodyweight change (from drug administration after 18hr fast to post-PR test refed state next day): *t*_(12)_=7.1, *P*<0.0001. **f**, latency to 1^st^ nose poke once test session initiated: *t*_(14)_=0.1389, *P=*0.8915. **g**, working time AUC (duration from first to last nose poke in each ratio, for rewards 1-10 in total): *t*_(15)_=3.3, *P*=0.0047. **h**, post-reinforcement pause AUC (duration from end of reward access to first nose poke for next ratio, for rewards 1-10 total): *t*_(15)_=2.2, *P*=0.0422. **i**, working time per reward: drug x reward # *F*_(9,135)_=13.23, *P*<0.0001. **j**, representative path plots tracking one mouse’s nose keypoint during the full PR task session under vehicle and sema conditions and when session length was aligned to the last matched reward (reward #7 in this example) earned under both conditions. **k**, distance travelled in full session: *t*_(10)_=9.7, *P*<0.0001. **l**, mean velocity during full session: *t*_(10)_=8.1, *P*<0.0001. **m**, distance travelled until last matched reward: *t*_(10)_=0.5, *P*=0.6221. **n**, time taken to obtain last matched reward: *t*_(10)_=4.3, *P*=0.0015. **o**, average velocity during time taken to obtain last matched reward: *t*_(10)_=5.0, *P*=0.0006. **p**, trajectories of manually annotated behavioural syllables from generated by unsupervised analysis using Keypoint MoSeq. **q**, PCA of syllable usage within the matched-reward window (circles show individual mice per session, triangles show group means) and pie charts of proportion of time observed for each behavioural grouping. PC1 and PC2 explained 47.7% and 11.2% of the variance, respectively. PC1 scores, vehicle vs semaglutide: *t*_(10)_=5.02, *P*=0.0005. **r**, immobility duration: *t*_(10)_=2.47, *P*=0.033. **s**, in-place activity duration: *t*_(10)_=3.62, *P*=0.0047. **t**, rearing duration: *t*_(10)_=4.54, *P*=0.0011. **u**, immobility %: *t*_(10)_=2.77, *P*=0.020. **v**, in-place activity %: *t*_(10)_=2.55, *P*=0.029. **w**, rearing %: *t*_(10)_=0.16, *P*=0.88.

**Extended Data Figure 2:**
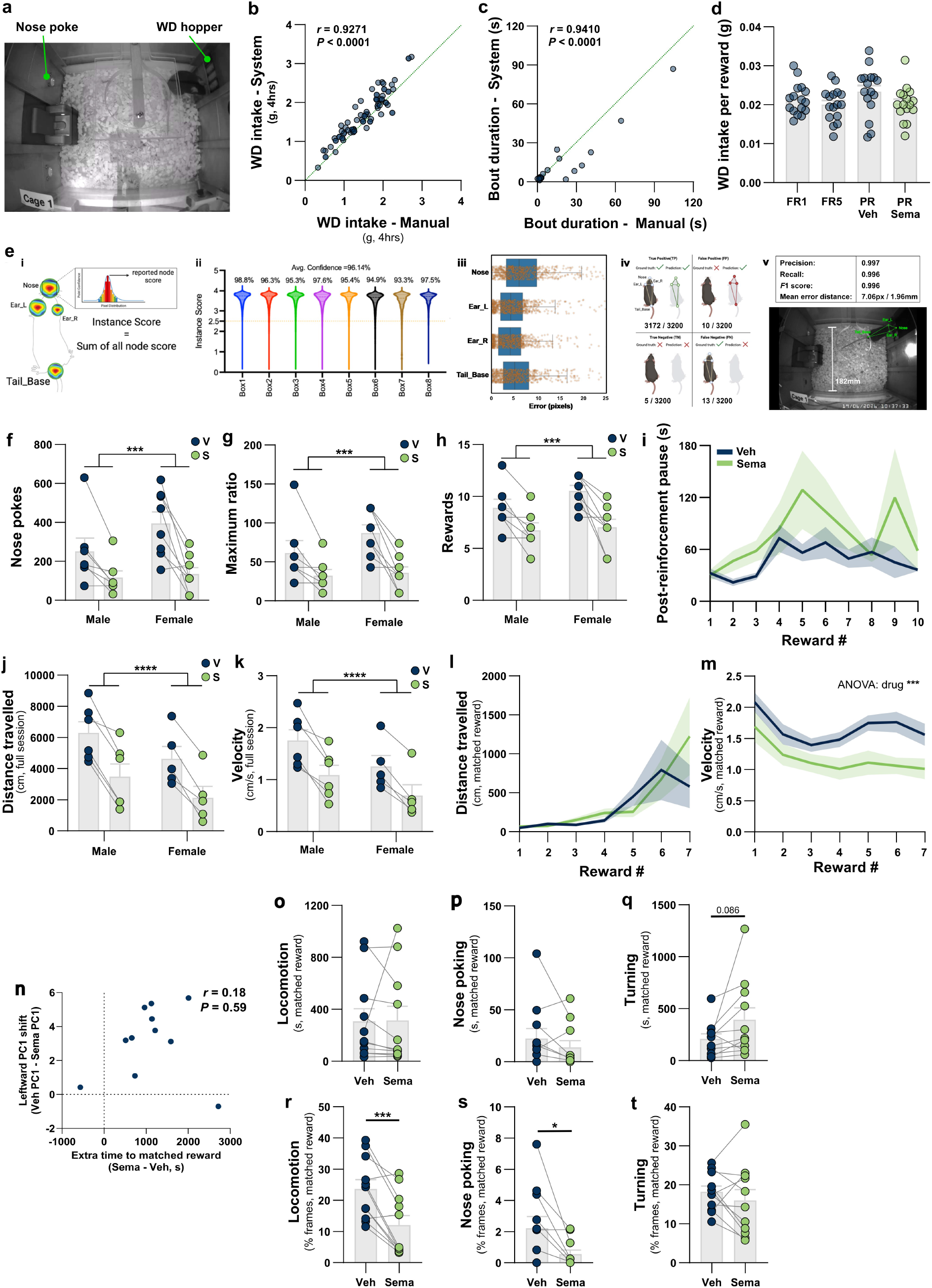
Semaglutide suppresses motivation to consume high-fat high-sugar Western diet. **a**, screenshot from top down camera view of custom eating behaviour analysis platform. **b**, validation of WD intake measurement using HM2++ system: correlation of mass of WD consumed as measured by monitoring system vs manual weighing of hopper after 4hr eating behaviour test, from 7 cages over 12 test sessions, Pearson *r*=0.9271, *P*<0.0001. **c**, validation of WD bout duration measurement using HM2++ system: correlation of WD bout duration as measured by monitoring system vs manual offline scoring of bout duration from video recordings, from 2 cages; Pearson *r*=0.9410, *P*<0.0001. **d**, WD intake consumed per 7s reward access period: reinforcement schedule *F*_(2.1, 31.2)_=2.7, *P*=0.0817. **e**, validation and performance of keypoint identification model developed for behavioural tracking during PR test: i) schematic of node-based pose estimation and instance score calculation; ii) trained model performance on novel mice across the 8 operant boxes; iii) model predicted node distances from manual ground truth annotations; iv) node visibility definitions and incidences used to derive performance metrics; v) model performance metrics, and example of keypoint identification on mouse, and conversion of pixel error to real-world distance in spatial context of operant box. **f**, total nose pokes by sex: drug *F*_(1,13)_=28.1, *P*=0.0001; drug x sex *F*_(1,13)_=2.8, *P*=0.1175. **g**, maximum ratio completed by sex: drug *F*_(1,13)_=29.9, *P*=0.0001; drug x sex *F*_(1,13)_=2.2, *P*=0.1583. **h**, total rewards earned by sex: drug *F*_(1,13)_=28.2, *P*=0.0001; drug x sex *F*_(1,13)_=1.63, *P*=0.2241. **i**, post-reinforcement pause per reward: drug *F*_(9,135)_=2.7, *P*=0.1194. **j**, distance travelled in full session by sex: drug *F*_(1,9)_=86.6, *P*<0.0001; drug x sex *F*_(1,9)_=0.3, *P*=0.5935. **k**, average velocity in full session by sex: drug *F*_(1,9)_=59.5, *P*<0.0001; drug x sex *F*_(1,9)_=0.5, *P*=0.5190. **l**, distance travelled over session (all mice to last matched reward earned, analysed up to reward # in which ≥50% of group have matched rewards): reward # *F*_(2.0,20.4)_=6.3, *P*=0.0074; drug *F*_(1.0,10.0)_=0.9, *P*=0.3744; drug x reward # *F*_(1.0,10.0)_=0.9, *P*=0.3744. **m,** average velocity over session (all mice to last matched reward earned, analysed up to reward # in which ≥50% of group have matched rewards): reward # *F*_(2.0,20.7)_=9.4, *P*=0.0011; drug *F*_(1.0,10.1)_=22.3, *P*=0.0008; drug x reward # *F*_(2.3,17.1)_=1.9, *P*=0.1779. **n**, PC1 shift vs additional time to matched reward for each mouse. Spearman *r*=0.18, *P*=0.59. **o**, locomotion duration: *t*_(10)_=0.08, *P*=0.94. **p,** nose poking duration: *t*_(10)_=1.3, *P*=0.22. **q**, turning duration: *t*_(10)_=1.91, *P*=0.086. **r**, locomotion %: *t*_(10)_=4.72, *P*=0.0008. **s,** nose poking %: *t*_(10)_=2.62, *P*=0.026. **t**, turning duration: *t*_(10)_=0.97, *P*=0.36.

We first used this system to investigate the effects of semaglutide on meal microstructure in intermittently WD-fed mice, during the 4hr active anorectic window we had previously identified (Fig. 1m). Semaglutide reduced 4hr WD intake by 51%, with no evidence of sex-dependence in the anorectic effect, and reduced 24hr bodyweight to a comparable degree as in chow-eating mice (Fig. 1n-p). Meal pattern analysis revealed that semaglutide potentiated both satiation and satiety in WD-feeding mice (Fig. 1q-t), consistent with its effects on chow-feeding mice in the metabolic phenotyping system. We then used a conditioned flavour avoidance paradigm (Fig. 1u), widely used and validated as a proxy for GLP-1RA-induced nausea / aversion^5,10,24,25^. Semaglutide elicited a potent avoidance response (Fig. 1v), consistent with negative valence in mice, and the high incidence of nausea and visceral malaise reported in humans^2^.

### Semaglutide suppresses motivation to consume high-fat high-sugar Western diet

In addition to the well-established effects of GLP-1RA drugs on the homeostatic processes of satiation and satiety, these drugs are also increasingly being investigated for their effects on hedonic behaviours, including on the rewarding effects of highly palatable food^3,26,27^. We therefore developed a progressive ratio (PR) reinforcement schedule operant task to test the effects of semaglutide on appetitive eating behaviours. Specifically, we quantified motivation to work for brief access to WD using an exponentially increasing schedule of reinforcement, designed to elicit a breakpoint before mice become satiated from consumption of rewards^28,29^. Motivation to consume WD was assessed using both standard breakpoint analysis, microstructural analysis of PR task performance using behavioural metrics derived from the operant response system, and from a machine-learning based video analysis pipeline for activity tracking and unsupervised behaviour identification (Fig. 2a, EDFig. 2e).

Following vehicle injection, mice on average completed 328 cumulative nose pokes and earned 10 rewards in a single session, with an average maximum ratio of 74 pokes for a single reward (Fig. 2b-d). Semaglutide reduced the number of nose pokes made and maximum ratio achieved by over 50%, such that mice earned fewer than 7 rewards on average (Fig. 2b-d). There was no evidence of sex differences in PR task performance (EDFig. 2f-h) and net bodyweight loss was similar to that under *ad libitum* chow- and WD-eating conditions (Fig. 2e).

To further investigate whether this reduced breakpoint reflected lower motivation rather than general behavioural suppression, we analysed the microstructure of task performance. Semaglutide did not affect the latency to initiate the task (Fig. 2f), but increased both working time (time taken to complete at individual ratio) and post-reinforcement pause duration (time taken to re-engage with the task after a reward) (Fig. 2g-i, ED Fig. 2i). Post-reinforcement pauses were lengthened throughout the session, possibly reflecting a reduction in incentive motivation^30,31^ (Fig. 2h), whereas increases in working time only emerged at higher response ratios (Fig. 2g-i).

We additionally tracked the locomotor activity of mice, to quantify distance travelled and average velocity during the PR assay. Semaglutide reduced distance and velocity by ∼50% and ∼40%, respectively, over the full session, in both male and female mice (Fig. 2j-l, EDFig. 2j-k). However, the reduced distance travelled over the full session simply reflected the earlier breakpoint, as this effect was lost when analysis was restricted to the period in which mice earned an equivalent number of rewards under both treatments (reward-matched analysis; Fig. 2m, EDFig. 2l). Semaglutide-treated mice did however take longer to earn the same number of rewards, lowering their average velocity, and as with working time, this slowing was greatest at higher ratios (Fig. 2n-o, EDFig. 2m). As behavioural testing occurred 2-3 h after injection, during the peak of acute homeostatic effects at this dose (Fig. 1b,n), effects emerging only at higher ratios are unlikely to reflect pharmacokinetic parameters of semaglutide. We extended the video analysis beyond tracking, using unsupervised machine-learning to identify behavioural syllables exhibited by mice during the PR assay. We manually curated these syllables into kinematically similar groupings (Fig. 2p) for analysis of semaglutide’s effects. Overall differences in behaviour were visualised by PCA, with PC1 and PC2 accounting for 47% and 11% of variance, respectively (Fig. 2q), and PC1 variance was not explained by slower task performance (Fig. 2n, EDFig. 2n). Instead, behavioural syllable variance was primarily driven by semaglutide-treated mice spending more time immobile, engaged in in-place activities (including grooming), and rearing (Fig. 2r-w). The proportion of time mice exhibited locomotion behaviours was consequently decreased, but the duration of these was unchanged, corroborating the tracking data. Nose poking was similarly only affected as a proportion but not absolute duration (as expected within matched reward windows), and turning behaviours were largely unaffected (EDFig. 2o-t). Together, these additional analyses of activity and behaviours during the PR assay reveal semaglutide-induced changes potentially consistent with increased off-task activity and/or visceral malaise. However, the specific changes at higher response requirements in particular suggest that generalised locomotor suppression is not driving reduced task performance, supporting the contention that semaglutide specifically reduces motivation to work for WD rewards.

Overall, these complementary metabolic and behavioural characterisations provide a comprehensive phenotypic profile of the acute effects of semaglutide on homeostatic and hedonic eating behaviours and whole-body metabolism, establishing a reference dataset for mechanistic studies to identify the neural circuit substrates of these phenotypic components.

### Semaglutide primarily recruits GLP-1R neurons in the AP and non-GLP-1R neurons in the NTS

GLP-1R-expressing neurons in the AP and NTS have been shown to suppress food intake by eliciting nausea and satiety, respectively, which has been concluded based on manipulation of the entire populations using *Glp1r*-Cre mouse lines or knockdown strategies^5,25^. However, several studies using various GLP-1RA drugs and labelled ligands have suggested that only a subset of DVC GLP-1R neurons are actually activated by these agonists, particularly in the NTS^8,9^, consistent with evidence that GLP-1RAs have limited access to this region^32–34^. This suggests that an unbiased, cell type-agnostic approach may be necessary to identify and functionally characterise the neural populations recruited by GLP-1RAs. We therefore employed an activity-dependent genetic labelling approach utilising the FosTRAP / TRAP2 transgenic mouse^35,36^ and using the previously characterised 60 μg/kg dose of semaglutide as the TRAPing stimulus, to reverse engineer the pathways which mediate semaglutide’s anorectic effects within the DVC (Fig. 3a). We first validated this approach using the FosTRAP mouse crossed with a tdTomato reporter strain, in which we determined that a common TRAPing protocol in which 4-hydroxy tamoxifen (4TM) is injected concurrently with the TRAPing stimulus^36,37^ was efficient but not selective for neurons activated by semaglutide vs vehicle TRAPed controls, particularly in the NTS (EDFig. 3b-c). We therefore employed an alternative protocol in which 4TM is given 3 hours after the stimulus^38^, and confirmed that with this protocol semaglutide specifically increased the number of TRAPed (Sema-TRAP) cells in both the AP and NTS compared to vehicle-injected mice (Veh-TRAP), that the number of TRAPed cells was comparable to those exhibiting semaglutide-induced cFos immunoreactivity, and that cre-dependent transgene expression required 4TM (EDFig. 3a,d-g). We also used this tdTomato reporter strain to validate the efficiency and specificity of Cre-dependent viral targeting of Sema-TRAP neurons in the NTS, using a cre-dependent eYFP reporter virus (EDFig. 3h-i). We further confirmed the reproducibility of sema-TRAP labelling, with the majority of virally-transduced TRAPed neurons in both the AP and NTS exhibiting cFos immunoreactivity following repeat semaglutide administration (EDFig. 3j-l). These validation experiments confirmed that our Sema-TRAP model provided an unbiased approach to neuroanatomically characterise semaglutide-recruited neuronal circuits in the DVC, with high selectivity over vehicle-injected controls, and enabled their viral transduction for *in vivo* interrogation of their contributions to the distinct phenotypic components of semaglutide’s action.

**Figure 3:**
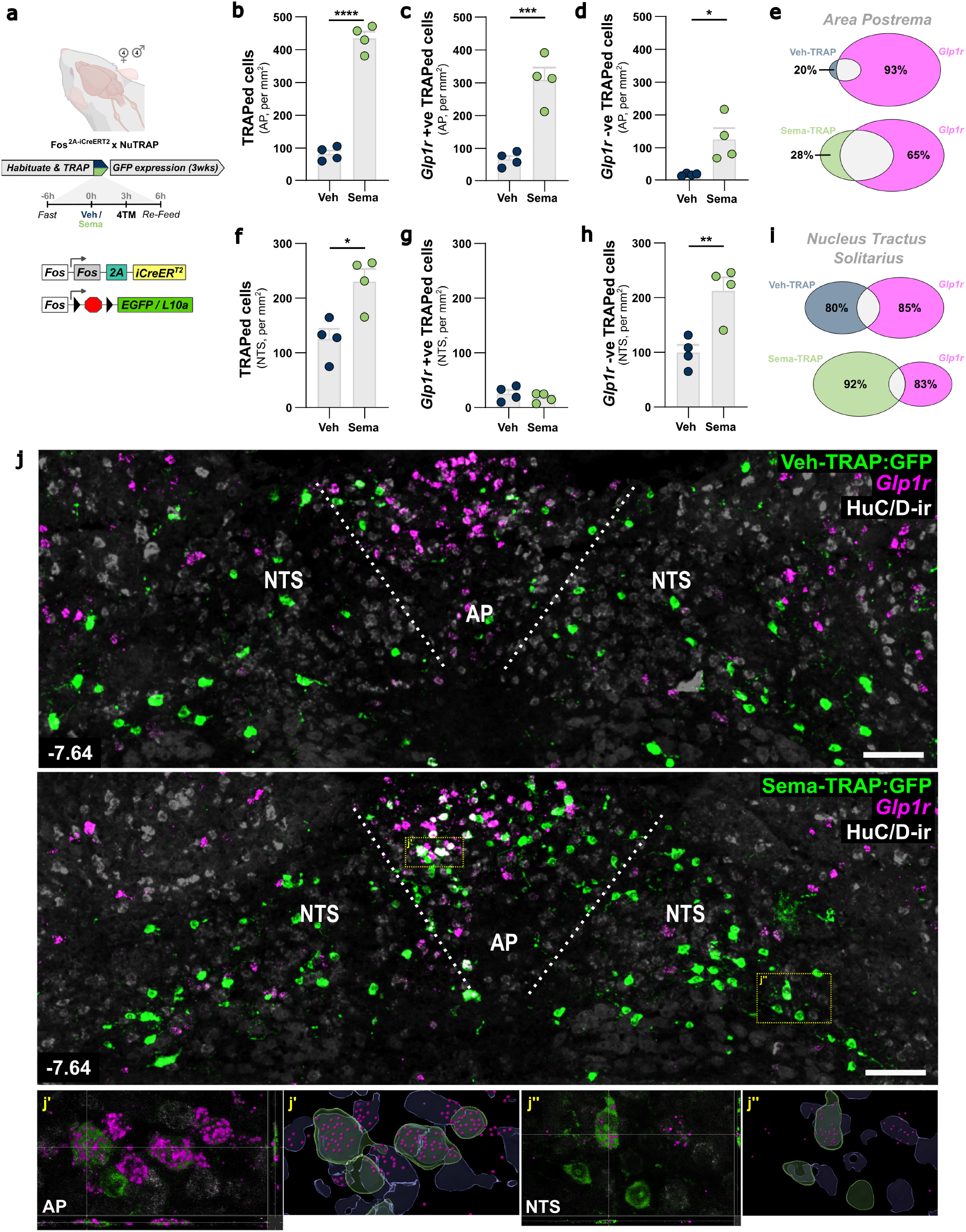
Semaglutide primarily recruits GLP-1R neurons in the AP and non-GLP-1R neurons in the NTS. **a** model schematic and experimental design for activity-dependent genetic labelling of semaglutide-activated neurons (Sema-TRAP). **b**, TRAPed cells in AP: *t*_(6)_=17.1, *P*<0.0001. **c**, *Glp1r* +ve TRAPed cells in AP: *t*_(6)_=5.8, *P*=0.0012. **d**, *Glp1r* -ve TRAPed cells in AP: *t*_(6)_=3.3, *P*=0.0155. **e**, proportion of *Glp1r* +ve and -ve cells in AP of Veh-TRAP and Sema-TRAP mice. **f**, TRAPed cells in NTS: *t*_(6)_=2.5, *P*=0.0490. **g**, *Glp1r* +ve TRAPed cells in NTS: *t*_(6)_=1.2, *P*=0.2870. **h**, *Glp1r* -ve TRAPed cells in NTS: *t*_(6)_=2.8, *P*=0.0311. **i**, proportion of *Glp1r* +ve and -ve cells in NTS of Veh-TRAP and Sema-TRAP mice. **j**, representative photomicrographs of coronal section of the dorsal vagal complex from Veh-TRAP and Sema-TRAP mice showing GFP and HuC/D immunoreactivity and *in situ* hybridisation for *Glp1r* mRNA, scale bars = 100μm. Insets show orthogonal projections and Imaris surfaces and spots 3D reconstructions to confirm colocalisation of GFP and/or HuC/D-ir soma and Glp1r puncta in AP (j’) and NTS (j’’).

**Extended Data Figure 3:**
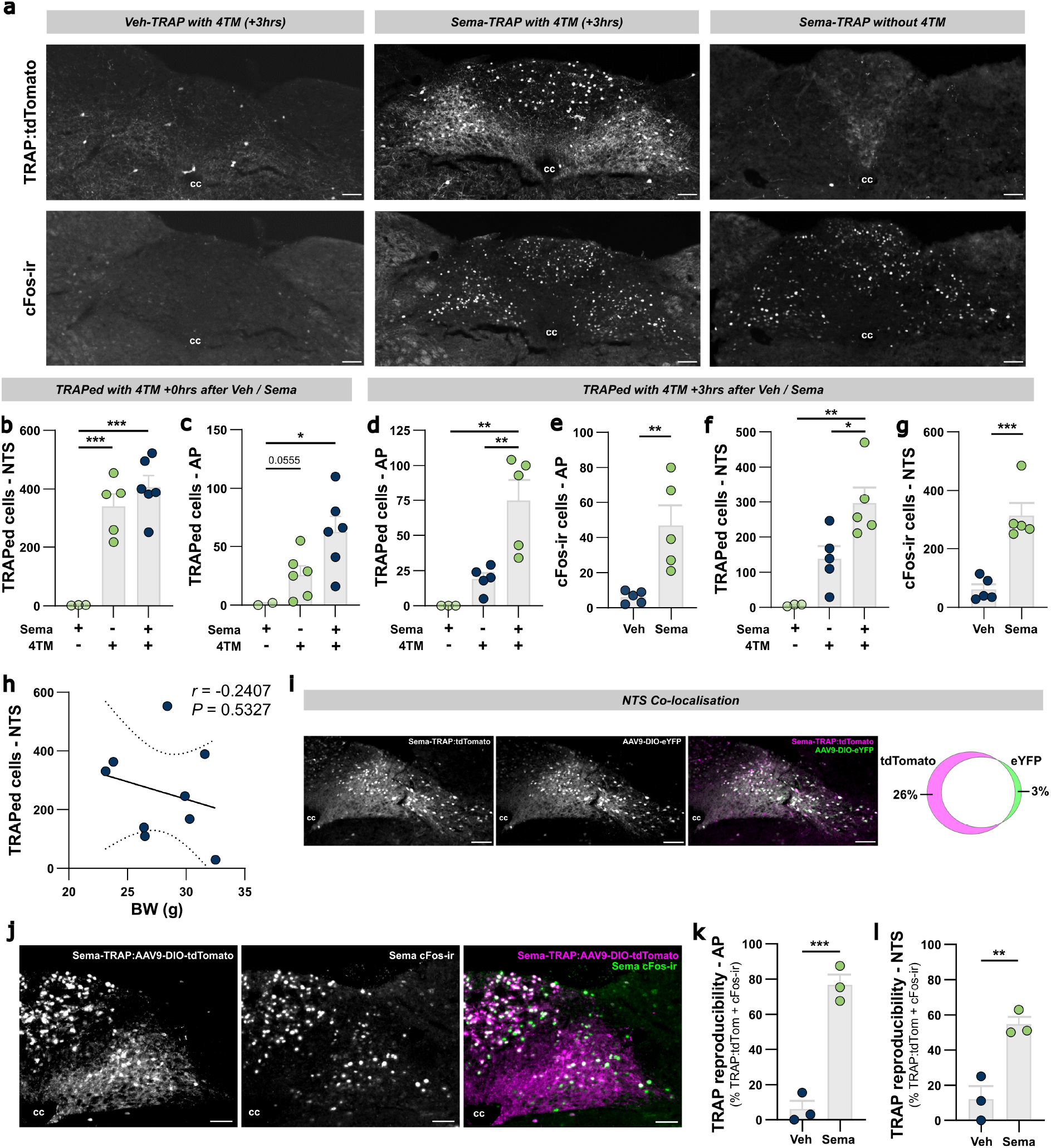
Optimisation and validation of the Sema-TRAP model. **a,** representative photomicrographs of DVC from Fos^2A-iCreERT2^:Ai9 mice administered vehicle or semaglutide as TRAPing stimulus, with or without 4-hydroxy tamoxifen (4TM) injected 3 hours after the TRAPing stimulus. DVC tissue was additionally processed for cFos immunoreactivity following a second administration of the same stimulus prior to perfuse-fixation. Scale bars = 100μm, cc: central canal. **b**, number of TRAPed cells in NTS (4TM +0hrs): drug *F*_(2,11)_=22.1, *P*=0.0001. **c**, number of TRAPed cells in AP (4TM +0hrs): drug *F*_(2,11)_=5.7, *P*=0.0199. **d**, number of TRAPed cells in AP (4TM +3hrs): drug *F*_(2,10)_=13.3, *P*=0.0015. **e**, number of cFos immunoreactive cells in AP: drug *t*_(4.6)_=13.3, *P*=0.0026. **f**, number of TRAPed cells in NTS: drug *F*_(2,10)_=12.1, *P*=0.0022. **g**, number of cFos immunoreactive cells in NTS: drug *t*_(3.5)_=13.3, *P*=0.0232. **h,** number of TRAPed cells in NTS of vehicle-TRAP control mice does not significantly correlate with mouse bodyweight at time of TRAPing, Pearson *r*=-0.2407, *P*=0.5327. **i**, representative photomicrographs of DVC and quantification summary from Sema-TRAP:Ai9 (tdTomato reporter) mice injected with AAV9-Ef1a-DIO-eYFP targeted to the NTS to validate the efficiency and specificity of viral transduction of TRAPed neurons in this region. Neurons with tdTomato and eYFP signals co-localised comprised 97% of all eYFP+ and 74% of all tdTomato+ neurons in the NTS (quantified from 6 coronal sections each from 3 mice). Scale bars = 100μm, cc: central canal. **j**, representative photomicrographs of DVC Sema-TRAP mice injected with AAV9-hSyn-DIO-tdTomato targeted to the AP + NTS to validate the reproducibility this model in virally-transduced DVC neuron populations, as assessed by quantification of the percentage of Sema-TRAP neurons expressing viral tdTomato which are also cFos immunoreactive when mice were given a repeated injection of the same TRAPing stimulus (vehicle or semaglutide) 3 weeks later, and then perfused 4 hours after drug administration. Scale bars = 100μm, cc: central canal. Quantification data obtained from 10 coronal sections from each of 3 mice per group. **k**, TRAP reproducibility in AP: drug *t*_(4)_=9.5, *P*=0.0007. **l**, TRAP reproducibility in NTS: drug *t*_(4)_=5.1, *P*=0.0069. All data presented as mean ± sem. * *P*<0.05, ** *P*<0.01, *** *P*<0.001.

**Extended Data Figure 4:**
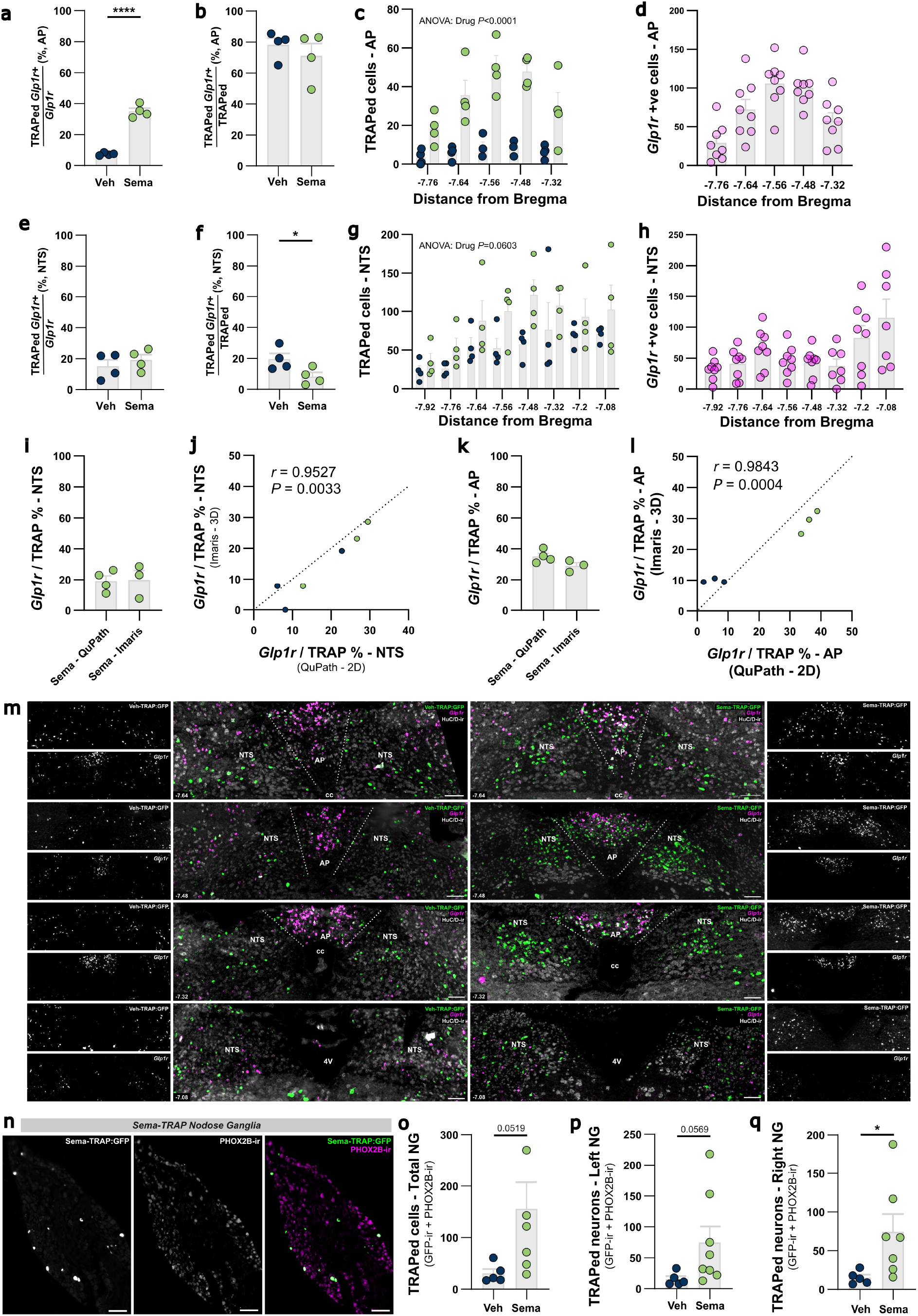
Semaglutide primarily recruits GLP-1R neurons in the AP and non-GLP-1R neurons in the NTS, and neurons in the nodose ganglia. **a**, % of *Glp1r* +ve cells in AP which were TRAPed: *t*_(6)_=13.2, *P*<0.0001. **b**, % of TRAPed cells in AP which were *Glp1r* +ve: *t*_(6)_=0.8, *P*=0.4763. **c**, number of TRAPed cells through rostro-caudal extent of the AP: drug *F*_(1,6)_=292.1 *P*<0.0001. **d**, number of *Glp1r* +ve cells through rostro-caudal extent of the AP. **e**, % of *Glp1r* +ve cells in NTS which were TRAPed: *t*_(6)_=2.5, *P*=0.0489. **f**, % of TRAPed cells in NTS which were *Glp1r* +ve: *t*_(6)_=2.6, *P*=0.0388. **g**, number of TRAPed cells through rostro-caudal extent of the NTS: drug *F*_(1,6)_=5.3 *P*=0.0603. **h**, number of *Glp1r* +ve cells through rostro-caudal extent of the AP. **i-j**, % of *Glp1r* +ve cells in NTS which were TRAPed (2D QuPath vs 3D Imaris pipelines): *t*_(5)_=0.1, *P*=0.9270, Pearson correlation statistics shown on plot. **k-l**, % of *Glp1r* +ve cells in AP which were TRAPed (2D QuPath vs 3D Imaris pipelines): *t*_(5)_=2.0, *P*=0.1012, Pearson correlation statistics shown on plot. **m**, representative photomicrographs of coronal sections (Bregma -7.64 to -7.08) of the dorsal vagal complex from Veh-TRAP and Sema-TRAP mice showing GFP and HuC/D immunoreactivity and *in situ* hybridisation for *Glp1r* mRNA, scale bars = 100μm. **n**, representative photomicrograph of nodose ganglia from Fos^2A-iCreERT2^:NuTRAP mice administered semaglutide as TRAPing stimulus, and processed for GFP and PHOX2B immunoreactivity, to specifically identify TRAPed vagal afferent neurons in the nodose ganglia (rather than PHOX2B -ve neurons in contiguous jugular ganglia). Scale bars = 100μm. **o**, total TRAPed cells in left + right NG: *t*_(6.3)_=2.4, *P*=0.0519. **p**, TRAPed cells in left NG: *t*_(7.5)_=2.3, *P*=0.0569. **q**, TRAPed cells in right NG: *t*_(6.3)_=2.6, *P*=0.0402.

We then utilised the FosTRAP mouse crossed with a Cre-dependent GFP reporter strain^39^ that efficiently labels the soma of TRAPed neurons even in fresh frozen tissue. This allowed us to use *in situ* hybridisation for *Glp1r* to conduct a high-sensitivity quantitative analysis of the extent to which semaglutide recruits both GLP-1R and non-GLP-1R neurons through the full rostro-caudal axis of the AP and visceral NTS (Fig. 3a,j, EDFig. 4m). Semaglutide recruited over 5-fold more neurons overall in the AP than in vehicle-TRAP controls, with the majority of these expressing *Glp1r* as expected, as well as an increased number of non-*Glp1r* neurons (Fig. 3b-e, EDFig. 4a-d). Semaglutide similarly increased recruitment of neurons in the NTS overall (Fig. 3f). Strikingly, however, this increase was driven by the recruitment of neurons which did not express *Glp1r* (Fig. 3g-i), with the proportion of NTS *Glp1r* neurons recruited not significantly increased (EDFig. 4e-h). Given the somewhat surprising nature of this result in the NTS, we corroborated our 2D imaging approach using 3D reconstruction of neuronal soma and labelled *Glp1r* puncta, which confirmed the accuracy of our original approach for quantifying the proportions of *Glp1r* and non-*Glp1r* neurons recruited by semaglutide (EDFig. 4i-l).

**Figure 4:**
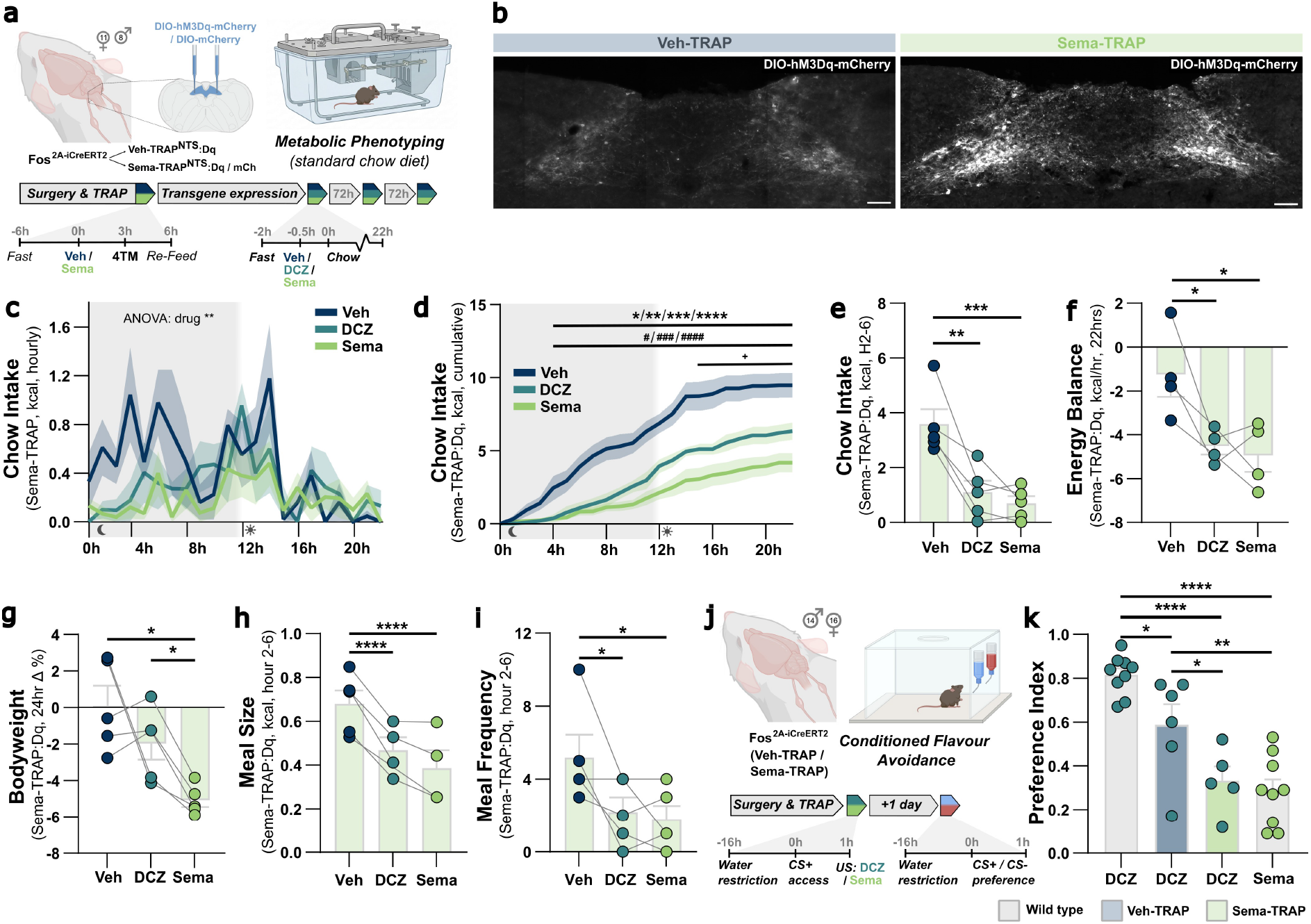
Semaglutide-recruited NTS neurons encode satiation and nausea. **a**, experimental model and design for metabolic phenotyping study in chow-fed veh-TRAP / Sema-TRAP mice transduced with cre-dependent hM3Dq-mCherry transgene in NTS. **b**, representative photomicrographs of hM3Dq-mCherry expression in the NTS of Veh-TRAP and Sema-TRAP mice, scale bars = 100μm. **c**, chow intake (Sema-TRAP:Dq, hourly non-cumulative): drug *F*_(1.0,4.2)_=31.4, *P*=0.0043. **d**, chow intake (Sema-TRAP:Dq, hourly cumulative): drug x hour *F*_(42,168)_=8.8, *P*<0.0001. **e**, chow intake (Sema-TRAP:Dq, hour 2-6): drug *F*_(2,8)_=21.5, *P*=0.0006. **f**, energy balance (Sema-TRAP:Dq, 22hrs): drug *F*_(2,6)_=11.7, *P*=0.0084. **g**, bodyweight (Sema-TRAP:Dq, 22hrs): drug *F*_(1.0,4.1)_=7.3, *P*=0.0529. **h**, meal size (Sema-TRAP:Dq, hour 2-6): drug *F*_(2,6)_=102.3, *P*<0.0001. **i**, meal frequency (Sema-TRAP:Dq, hour 2-6): drug *F*_(1.7,6.7)_=7.3, *P*=0.0235. **j**, experimental model and design for conditioned flavour avoidance assay in veh-TRAP / Sema-TRAP mice. **k**, preference index in CFA: drug *F*_(3,25)_=19.8, *P*<0.0001. All data presented as mean ± sem. Pairwise comparisons on timecourse plots: * Veh vs Sema, # Veh vs DCZ, + DCZ vs Sema. */#/+ *P*<0.05, **/##/++ *P*<0.01, ***/###/+++ *P*<0.001, ****/####/++++ *P*<0.0001.

**Extended Data Figure 5:**
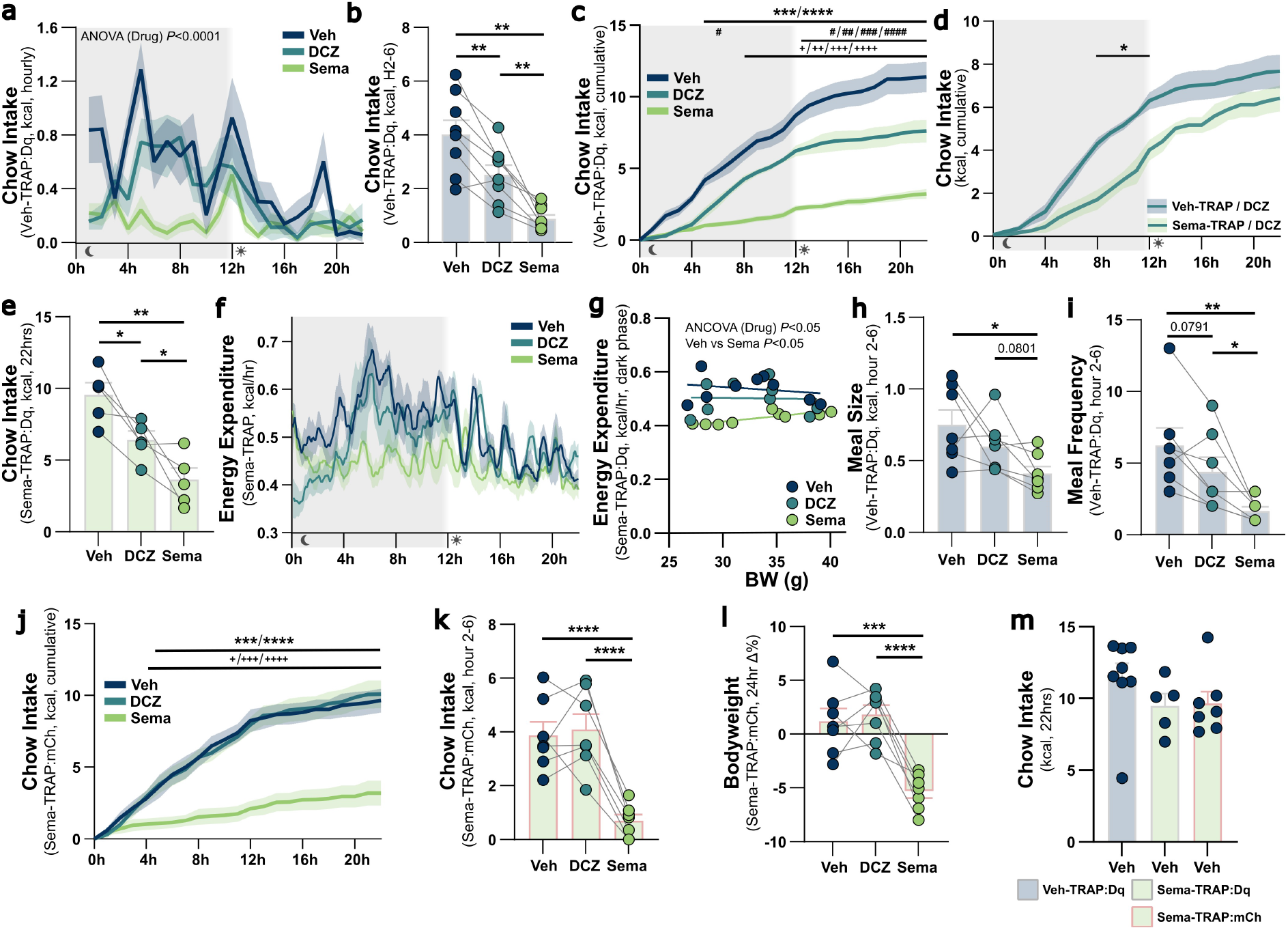
Semaglutide-recruited NTS neurons encode satiation and nausea. **a,** chow intake (Veh-TRAP:Dq, hourly non-cumulative): drug *F*_(1.8,12.5)_=33.5, *P*<0.0001. **b,** chow intake (Veh-TRAP:Dq, hour 2-6): drug *F*_(1.4,9.8)_=27.2, *P*=0.0002. **c,** chow intake (Veh-TRAP:Dq, hourly cumulative): drug x hour *F*_(42,294)_=15.7, *P*<0.0001. **d,** chow intake (Veh-TRAP:Dq + Sema-TRAP:Dq, hourly cumulative): drug x hour *F*_(21,231)_=1.6, *P*=0.0432. **e,** chow intake (Sema-TRAP:Dq, 22hrs): drug *F*_(2,8)_=23.4, *P*=0.0005. **f,** energy expenditure (Sema-TRAP:Dq) plotted as 5 minute bins with 30 minute rolling average (for visualisation purposes only, unsmoothed hourly data used for ANCOVA analysis. **g**, energy expenditure (Sema-TRAP:Dq, dark phase) ANCOVA: drug *F*_(2,11)_=4.3, *P*=0.0409, Veh vs Sema *P*=0.0414. **h**, meal size (Veh-TRAP:Dq, hour 2-6): drug *F*_(1.3,7.6)_=6.7, *P*=0.0290. **i**, meal frequency (Veh-TRAP:Dq, hour 2-6): drug *F*_(2,12)_=11.3, *P*=0.0017. **j**, chow intake (Sema-TRAP:mCherry, hourly cumulative): drug x hour *F*_(42,252)_=21.5, *P*<0.0001. **k**, chow intake (Sema-TRAP:mCherry, hour 2-6): drug *F*_(2,12)_=35.8, *P*<0.0001. **l**, 24hr bodyweight change (Sema-TRAP:mCherry): drug *F*_(2,12)_=26.6, *P*<0.0001. **k**, chow intake (Veh + Sema-TRAP:Dq/mCh, 22hrs): drug *F*_(2,17)_=1.2, *P*=0.3141. All data presented as mean ± sem. Pairwise comparisons on timecourse plots: * Veh vs Sema, # Veh vs DCZ, + DCZ vs Sema. */#/+ *P*<0.05, **/##/+ + *P*<0.01, ***/###/+++ *P*<0.001, ****/####/++++ *P*<0.0001.

Given our NTS TRAPing results strongly implicate that the majority of this activation must be indirect, driven by inputs from alternative, more accessible sites of action, we also investigated whether semaglutide recruited vagal afferent neurons, by quantifying GFP expression in PHOX2B-immunoreactive vagal neurons. PHOX2B expression identifies nodose ganglia neurons specifically, distinguishing them from the contiguous jugular neuron population^40^. Interestingly, despite previous evidence that vagal signalling is not necessary for the full anorectic or weight loss effects of GLP-1RA drugs^5,34^, we observed a recruitment of vagal afferent neurons by semaglutide, with similar activation levels observed in both left and right nodose ganglia (EDFig. 4n-q).

These data demonstrate that semaglutide activates a substantial population of GLP-1R-expressing neurons in the AP, consistent with this being a direct site of action. However, the majority of the NTS neuron population(s) recruited by semaglutide do not express the receptor upon which this drug acts. This implies that they are downstream of Sema-TRAP neurons in the AP and/or other sites of action, perhaps including GLP-1R-expressing vagal afferent neurons. These semaglutide-recruited NTS neurons are therefore likely distinct from the NTS GLP-1R population shown to be sufficient for non-aversive satiety^5^, supporting the rationale that a cell type-agnostic approach is required to functionally interrogate which phenotypic components they mediate.

### Semaglutide-recruited NTS neurons encode satiation and nausea

To reverse engineer the phenotypic roles of semaglutide-recruited circuits in the NTS, we first transduced Sema-TRAP^NTS^ neurons with the activating DREADD hM3Dq. We then reactivated these Sema-TRAP^NTS^:Dq neurons in mice undergoing metabolic and behavioural phenotyping to compare the effects of DCZ-induced reactivation to both vehicle, and semaglutide itself (Fig. 4a-b).

Chemogenetic reactivation of Sema-TRAP^NTS^:Dq neurons suppressed chow intake to a comparable degree as semaglutide over the first half of the dark phase, including during the hour 2-6 acute semaglutide activity window, with cumulative intake only diverging in the light phase (Fig. 4c-e). Consistent with the small population of stochastically active Veh-TRAP neurons seen in the NTS (Fig. 3f), reactivation of Veh-TRAP^NTS^:Dq neurons elicited a modest suppression of intake, that was significantly weaker than either semaglutide itself or reactivation of Sema-TRAP^NTS^:Dq neurons (EDFig. 5a-d).

**Figure 5:**
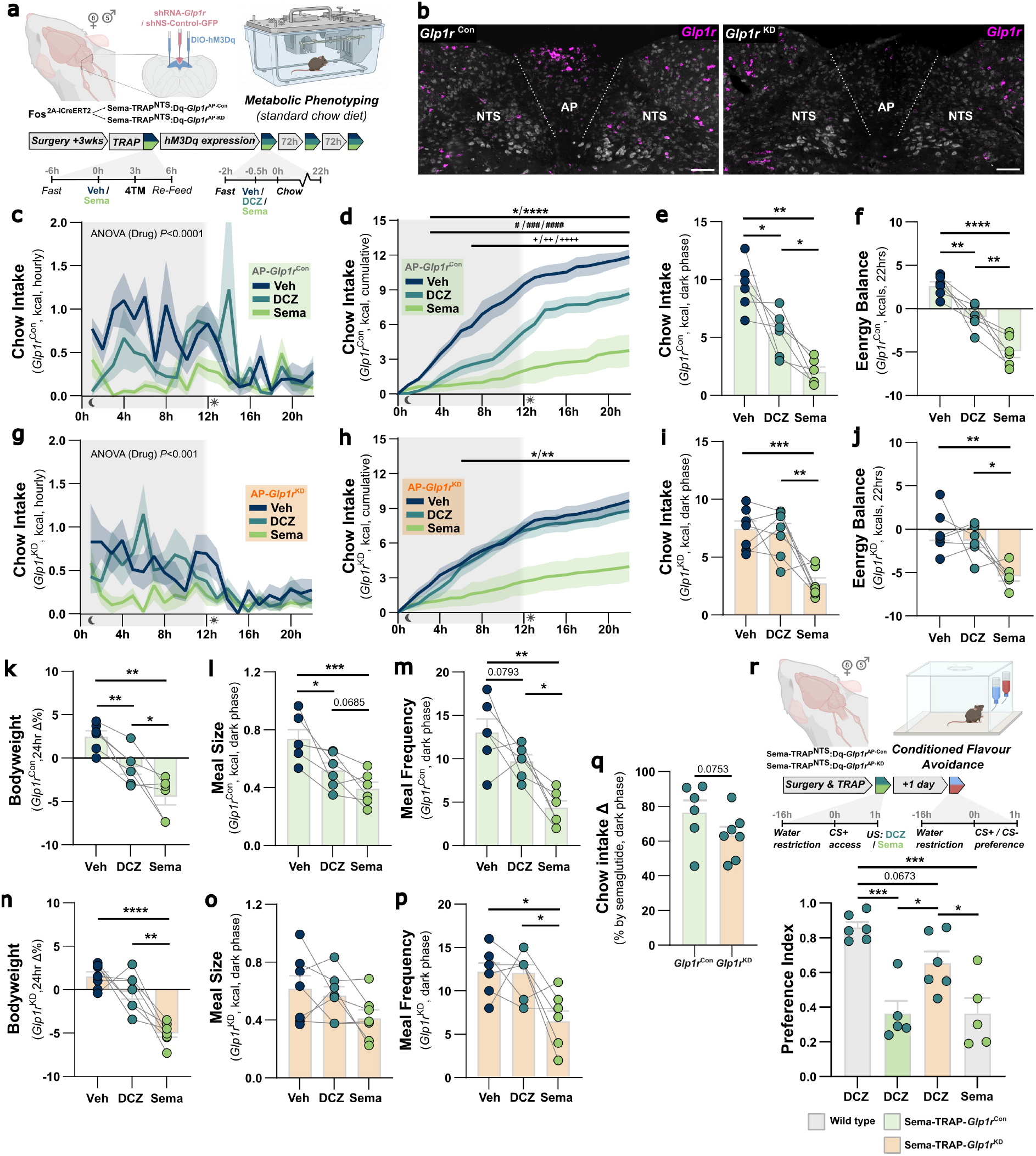
Semaglutide-induced satiation and nausea are mediated by area postrema GLP-1R. **a**, experimental model and design for metabolic phenotyping study in chow-fed mice transduced with cre-dependent hM3Dq in NTS and Sema-TRAPed 3 weeks after shRNA-mediated knockdown of AP *Glp1r (Glp1r*^KD^), or injected with non-silencing shRNA-GFP control virus (*Glp1r*^Con^). **b**, representative photomicrographs showing intact Glp1r RNA in *Glp1r*^Con^ controls mice, and AP-selective knockdown in *Glp1r*^KD^ mice. **c**, chow intake (Sema-TRAP^NTS^:Dq-*Glp1r* ^AP-Con^, hourly non-cumulative): drug *F*_(1.7,8.6)_=42.0, *P*<0.0001. **d**, chow intake (Sema-TRAP^NTS^:Dq-*Glp1r* ^AP-Con^, hourly cumulative): drug x hour *F*_(42,210)_=14.7, *P*<0.0001. **e**, chow intake (Sema-TRAP^NTS^:Dq-*Glp1r* ^AP-Con^, dark phase): drug *F*_(1.5,7.5)_=19.1, *P*=0.0017. **f**, energy balance (Sema-TRAP^NTS^:Dq-*Glp1r* ^AP-Con^, 22hrs): drug *F*_(2,10)_=34.1, *P*<0.0001. **g**, chow intake (Sema-TRAP^NTS^:Dq-*Glp1r* ^AP-KD^, hourly non-cumulative): drug *F*_(1.5,9.2)_=19.6, *P*=0.0008. **h**, chow intake (Sema-TRAP^NTS^:Dq-*Glp1r* ^AP-KD^, hourly cumulative): drug x hour *F*_(3.0,17.8)_=13.9, *P*<0.0001. **i**, chow intake (Sema-TRAP^NTS^:Dq-*Glp1r* ^AP-KD^, dark phase): drug *F*_(2,12)_=15.4, *P*=0.0005. **j**, energy balance (Sema-TRAP^NTS^:Dq-*Glp1r* ^AP-KD^, 22hrs): drug *F*_(1.5,7.5)_=8.9, *P*=0.0136. **k**, bodyweight (Sema-TRAP^NTS^:Dq-*Glp1r* ^AP-Con^, 24hrs): drug *F*_(1.4,7.2)_=23.5, *P*=0.0010. **l**, meal size (Sema-TRAP^NTS^:Dq-*Glp1r* ^AP-Con^, dark phase): drug *F*_(2,10)_=15.3, *P*=0.0009. **m**, meal frequency (Sema-TRAP^NTS^:Dq-*Glp1r* ^AP-Con^, dark phase): drug *F*_(2,10)_=13.3, *P*=0.0016. **n**, bodyweight (Sema-TRAP^NTS^:Dq-*Glp1r* ^AP-KD^, 24hrs): drug *F*_(1.2,7.0)_=23.7, *P*=0.0015. **o**, meal size (Sema-TRAP^NTS^:Dq-*Glp1r* ^AP-KD^, dark phase): drug *F*_(2,12)_=3.1, *P*=0.0848. **p**, meal frequency (Sema-TRAP^NTS^:Dq-*Glp1r* ^AP-KD^, dark phase): drug *F*_(2,12)_=7.8, *P*=0.0067. **q**, % reduction in dark phase chow intake induced by semaglutide in Sema-TRAP^NTS^:Dq-*Glp1r* ^AP-Con^ vs Sema-TRAP^NTS^:Dq-*Glp1r* ^AP-KD^ mice: *t*_(11)_=1.55, *P*=0.0753 (one-tailed). **r,** experimental model and design for conditioned flavour avoidance assay in Sema-TRAP^NTS^:Dq-*Glp1r* ^AP-Con^ and Sema-TRAP^NTS^:Dq-*Glp1r*^AP-KD^ mice, preference index in CFA: drug *F*_(3,18)_=13.4, *P*<0.0001.

**Extended Data Figure 6:**
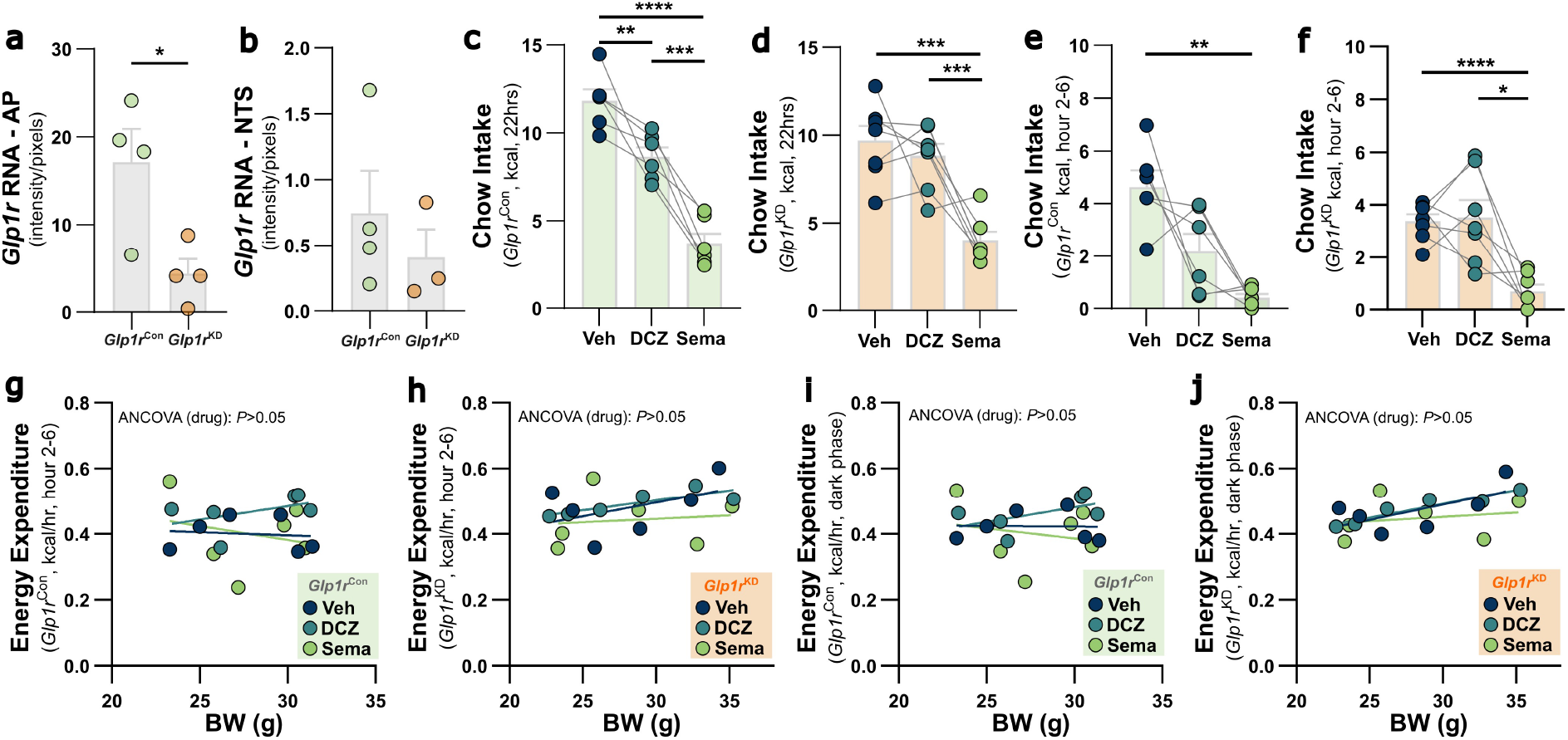
Semaglutide-induced satiation and nausea are mediated by area postrema GLP-1R. **a,** *Glp1r* knockdown quantification in AP: *t*_(6)_=2.7, *P*=0.0343. **b**, *Glp1r* knockdown quantification in NTS: *t*_(5)_=0.8, *P*=0.4569. **c**, chow intake (Sema-TRAP^NTS^:Dq-*Glp1r* ^AP-Con^, 22hrs): drug *F*_(2,10)_=42.0, *P*<0.0001. **d**, chow intake (Sema-TRAP^NTS^:Dq-*Glp1r* ^AP-KD^, 22hrs): drug *F*_(2,12)_=19.6, *P*=0.0002. **e**, chow intake (Sema-TRAP^NTS^:Dq-*Glp1r* ^AP-Con^, hour 2-6): drug *F*_(1.1,5.4)_=11.0, *P*=0.0181. **e**, chow intake (Sema-TRAP^NTS^:Dq-*Glp1r* ^AP-KD^, hour 2-6): drug *F*_(1.1,6.5)_=13.4, *P*=0.0087. **g**, energy expenditure (Sema-TRAP^NTS^:Dq-*Glp1r* ^AP-Con^, hour 2-6): drug *F*_(2,14)_=1.4, *P*=0.2831. **h**, energy expenditure (Sema-TRAP^NTS^:Dq-*Glp1r* ^AP-KD^, hour 2-6): drug *F*_(2,17)_=0.1, *P*=0.9101. **i**, energy expenditure (Sema-TRAP^NTS^:Dq-*Glp1r* ^AP-Con^, dark phase): drug *F*_(2,14)_=1.2, *P*=0.3311. **j**, energy expenditure (Sema-TRAP^NTS^:Dq-*Glp1r* ^AP-KD^, dark phase): drug *F*_(2,17)_=0.04, *P*=0.9597.

Reactivation of Sema-TRAP^NTS^:Dq neurons was sufficient to suppress 22 hr cumulative intake, but to a substantially lesser extent than semaglutide itself (EDFig. 5e). While semaglutide decreased dark phase energy expenditure, consistent with our earlier experiment (EDFig. 5f-g, EDFig. 1b), there was notably no effect of Sema-TRAP^NTS^:Dq neuron reactivation, potentially indicating this effect of semaglutide is mediated by site(s) of action outside of the DVC. The energy balance deficit over 22hrs after reactivation was similar to that after semaglutide, presumably due to the absence of an EE reduction being equivalent to the lower reduction in energy intake in DCZ vs semaglutide. Mice did not exhibit significant bodyweight loss following reactivation in this experiment (Fig. 4f-g), however this is likely an artefact of within-group variability, given the clear effect seen from the same manipulation in our subsequent experiment (Fig. 5g). The acute anorectic effect of Sema-TRAP^NTS^:Dq neuron reactivation was driven by reductions in both meal size and frequency, both of which were comparable to the effects of semaglutide during the acute activity window (Fig. 4h-i). This was in notable contrast to the negligible effects of Veh-TRAP^NTS^:Dq reactivation on meal size and modest effect on meal frequency over the same time period, neither of which approached the magnitude of semaglutide’s effect on these meal pattern parameters (EDFig. 5h-i).

Importantly, to control for non-specific effects of DCZ administration and Cre-dependent viral transduction, we labelled Sema-TRAP^NTS^ neurons with a Cre-dependent mCherry reporter virus. In these control mice, DCZ had no effect on cumulative food intake, hour 2-6 chow intake or bodyweight, whereas semaglutide produced effects consistent with those observed in the other cohorts (EDFig. 5j-l). Furthermore, 22hr food intake following vehicle injection was consistent between hM3Dq- and mCherry-transduced controls (EDFig. 5m). This demonstrates that the effects on eating behaviour in the Sema-TRAP^NTS^:Dq model were not confounded by off-target effects of DCZ, or constitutive effects of hM3Dq expression.

We then tested whether reactivation of Veh-TRAP^NTS^:Dq and Sema-TRAP^NTS^:Dq neurons was sufficient to condition an avoidance response, and how it compared to that elicited by semaglutide (Fig. 4j). Veh-TRAP^NTS^:Dq reactivation produced a modest reduction in CS preference, likely driven by TRAPing of a small population of NTS neurons active during the labelling window, consistent with the stochastic Veh-TRAP labelling. Reactivation of Sema-TRAP^NTS^:Dq neurons elicited a significant avoidance of the CS, reducing preference to 33%, comparable in magnitude to semaglutide itself (Fig. 4k).

These results demonstrate that the largely non-GLP-1R neuronal population(s) in the NTS which are activated by semaglutide are capable of suppressing food intake by advancing satiation and satiety, and can elicit a negative valence state consistent with the nausea-inducing effect of semaglutide.

### Semaglutide-induced satiation and nausea are mediated by area postrema GLP-1R

Our results show that chemogenetic reactivation of Sema-TRAP^NTS^ neurons is sufficient to potentiate satiation and elicit an avoidance response. This differs from previous reports that activation of NTS GLP-1R-expressing neurons suppresses food intake by non-aversive satiety^5^, but is consistent with our finding that semaglutide recruits a largely non-GLP-1R NTS population. Instead, the effects we observed following activation this non-GLP-1R population closely resemble the effects of activating AP GLP-1R neurons reported by others^5,25^. We therefore reasoned that Sema-TRAP^NTS^ neurons are likely downstream of AP GLP-1R neurons, and comprise an indirectly activated component of the same circuit(s) recruited by semaglutide to elicit satiation and nausea. We tested this hypothesis by selectively knocking down expression of *Glp1r* in the AP, prior to performing the same Sema-TRAP procedure in mice transduced with Cre-dependent hM3Dq in the NTS (Fig. 5a). This approach allowed us to isolate the phenotypic contributions which require AP GLP-1R, by directly comparing the effects of chemogenetic reactivation of the remaining NTS population that was still Sema-TRAPed despite knockdown of these receptors (Sema-TRAP^NTS^:Dq-*Glp1r*^AP-KD^) to a control group with intact AP GLP-1R (Sema-TRAP^NTS^:Dq-*Glp1r*^AP-Con^). Our knockdown strategy utilised virally-delivered shRNA, targeted to maximise transduction efficiency within the AP while minimising spread to adjacent regions, achieved ∼75% reduction in *Glp1r* signal intensity within the AP, while leaving *Glp1r* expression in the NTS intact (Fig. 5b, EDFig. 6a-b).

**Figure 6:**
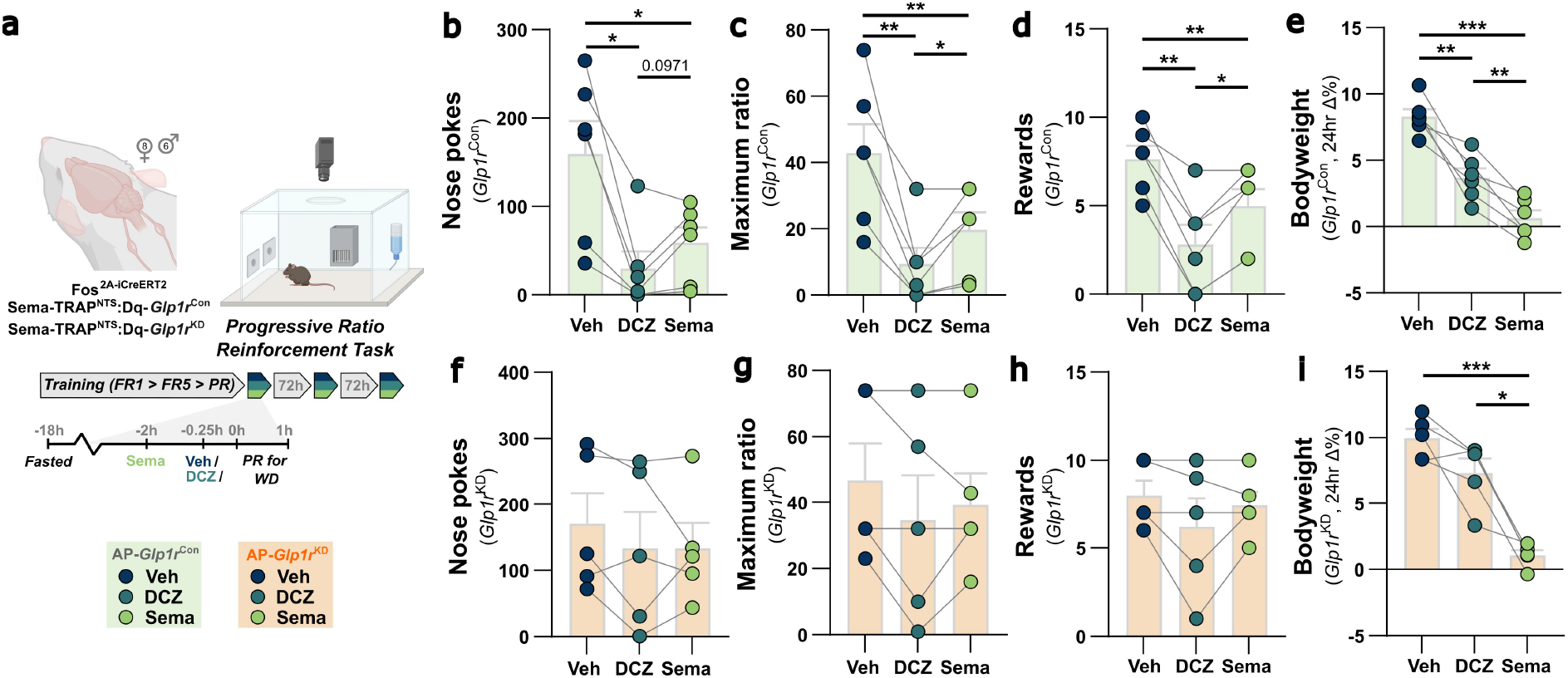
Semaglutide-induced suppression of motivation to consume Western diet requires area postrema GLP-1R. **a**, experimental model and design for progressive ratio (PR) schedule of reinforcement task for WD rewards in mice transduced with cre-dependent hM3Dq in NTS and Sema-TRAPed after knockdown of AP *Glp1r (Glp1r*^KD^), or injected with scrambled-GFP control virus in AP (*Glp1r*^Con^). **b**, total nose pokes (Sema-TRAP^NTS^:Dq-*Glp1r* ^AP-Con^): drug *F*_(1.8,5.9)_=17.1, *P*=0.0055. **c**, maximum ratio achieved (Sema-TRAP^NTS^:Dq-*Glp1r* ^AP-Con^): drug *F*_(1.2,5.9)_=22.1, *P*=0.0029. **d**, total reward earned (Sema-TRAP^NTS^:Dq-*Glp1r* ^AP-Con^): drug *F*_(1.4,7.0)_=42.2, *P*=0.0002. **e**, 24hr bodyweight change (from drug administration after 18hr fast to post-PR test refed state next day, Sema-TRAP^NTS^:Dq-*Glp1r* ^AP-Con^): drug *F*_(1.5,7.3)_=53.1, *P*<0.0001. **f**, total nose pokes (Sema-TRAP^NTS^:Dq-*Glp1r* ^AP-KD^): drug *F*_(2,8)_=1.1, *P*=0.3923. **g**, maximum ratio achieved (Sema-TRAP^NTS^:Dq-*Glp1r* ^AP-KD^): drug *F*_(2,8)_=2.2, *P*=0.1688. **h**, total reward earned (Sema-TRAP^NTS^:Dq-*Glp1r* ^AP-KD^): drug *F*_(1.2,5.0)_=2.5, *P*=0.1784. **i**, 24hr bodyweight change (from drug administration after 18hr fast to post-PR test refed state next day, Sema-TRAP ^NTS^:Dq-*Glp1r* ^AP-KD^): drug *F*_(1.4,5.5)_=36.2, *P*=0.0010.

**Extended Data Figure 7:**
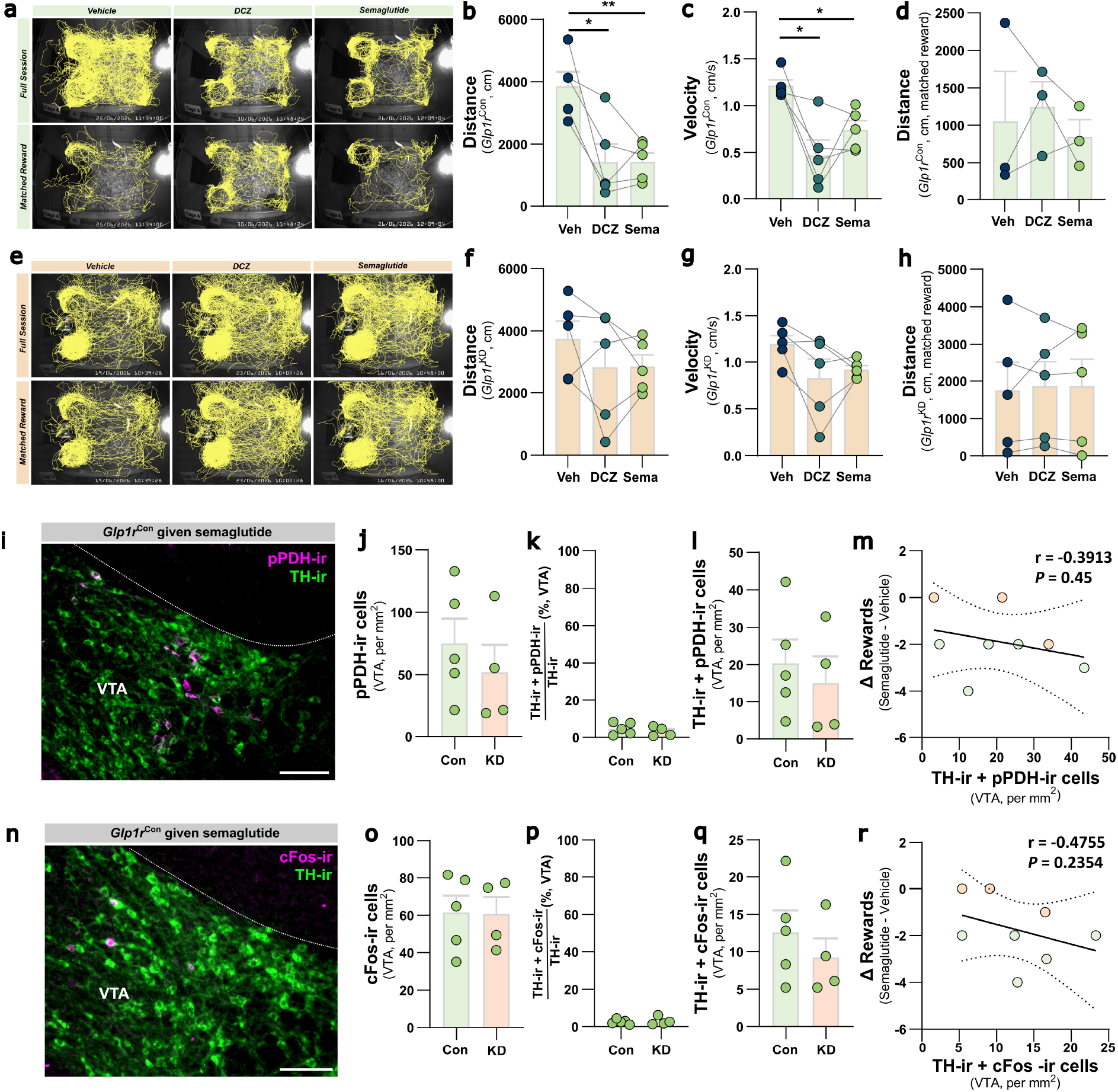
Semaglutide-induced suppression of motivation to consume Western diet is mediated by area postrema GLP-1R. **a**, representative path plots tracking one Sema-TRAP^NTS^:Dq-*Glp1r* ^AP-Con^ mouse’s nose keypoint during the full PR task session under vehicle, DCZ and sema conditions and when session length was aligned to the last matched reward earned under all conditions (reward #4 in this example). **b**, distance travelled in full session (Sema-TRAP^NTS^:Dq-*Glp1r* ^AP-Con^): drug *F*_(1.1,4.4)_=16.9, *P*=0.0117. **c**, mean velocity in full session (Sema-TRAP^NTS^:Dq-*Glp1r* ^AP-Con^): drug *F*_(1.7,6.9)_=9.8, *P*=0.0108. **d**, distance travelled until last matched reward earned (Sema-TRAP^NTS^:Dq-*Glp1r* ^AP-Con^): drug *F*_(1.7,3.5)_=0.4, *P*=0.6530. **e**, representative path plots tracking one Sema-TRAP^NTS^:Dq-*Glp1r* ^AP-KD^ mouse’s nose keypoint during the full PR task session under vehicle, DCZ and sema conditions and when session length was aligned to the last matched reward earned under all conditions (reward #7 in this example). **f**, distance travelled in full session (Sema-TRAP^NTS^:Dq-*Glp1r* ^AP-KD^): drug *F*_(1.8,7.1)_=1.1, *P*=0.3684. **g**, mean velocity in full session (Sema-TRAP^NTS^:Dq-*Glp1r* ^AP-KD^): drug *F*_(1.3,5.1)_=3.7, *P*=0.1100. **h**, distance travelled until last matched reward earned (Sema-TRAP^NTS^:Dq-*Glp1r* ^AP-KD^): drug *F*_(1.1,4.2)_=0.1, *P*=0.8030. **i**, representative photomicrograph of phosphorylated pyruvate dehydrogenase immunoreactivity (pPDH-ir) and tyrosine hydroxylase immunoreactivity (TH-ir) in the ventral tegmental area (VTA) of Sema-TRAP^NTS^:Dq-*Glp1r* ^AP-Con^ mouse administered semaglutide prior to transcardial perfusion, scale bar 100μm. **j**, number of pPDH-ir cells in the VTA: *t*_(7)_=0.77, *P*=0.4658. **k**, percentage of VTA TH-ir cells which are also pPDH-ir: *t*_(7)_=0.80, *P*=0.4505. **l**, number of VTA TH-ir cells which are also pPDH-ir: *t*_(7)_=0.55, *P*=0.5993. **l**, correlation between effect of semaglutide on number of rewards earned in PR task and number of VTA TH-ir cells which are also pPDH-ir: Spearman’s *r*=-0.3913, *P*=0.4500. **n**, representative photomicrograph of cFos immunoreactivity (cFos-ir) and tyrosine hydroxylase immunoreactivity (TH-ir) in the ventral tegmental area (VTA) of Sema-TRAP^NTS^:Dq-*Glp1r* ^AP-Con^ mouse administered semaglutide prior to transcardial perfusion, scale bar 100μm. **o**, number of cFos-ir cells in the VTA: *t*_(7)_=0.06, *P*=0.9566. **p**, percentage of VTA TH-ir cells which are also cFos-ir: *t*_(7)_=0.32, *P*=0.7551. **q**, number of VTA TH-ir cells which are also cFos-ir: *t*_(7)_=0.85, *P*=0.4266. **r**, correlation between effect of semaglutide on number of rewards earned in PR task and number of VTA TH-ir cells which are also cFos-ir: Spearman’s *r*=-0.4755, *P*=0.2354.

Reactivation of Sema-TRAP^NTS^:Dq-*Glp1r*^AP-Con^ neurons suppressed food intake over the circadian cycle relative to vehicle, but to a lesser extent than semaglutide itself (Fig. 5c-d, EDFig. 6c). This reactivation effect was also observed during the hour 2-6 window (EDFig. 6e) and the full dark phase (Fig. 5e), with a clearer effect during this longer period. Strikingly, reactivation of Sema-TRAP^NTS^:Dq-*Glp1r*^AP-KD^ neurons had no effect on cumulative intake over the entire circadian cycle, dark phase, or hour 2-6, but an anorectic effect of semaglutide itself was preserved (Fig. 5g-i, EDFig. 6d,f). There was no evidence of effects on energy expenditure during these windows (EDFig. 6g-j). Mice were in a negative energy balance 22hrs after both reactivation of Sema-TRAP^NTS^:Dq-*Glp1r*^AP-Con^ neurons and semaglutide, with a greater effect in the latter (Fig. 5f), which elicited an equivalent pattern of bodyweight loss (Fig. 5k). Conversely, in mice with AP GLP-1R expression knocked down, reactivation had no effect on energy balance or bodyweight, but semaglutide-induced loss was comparable to that seen in control mice (Fig. 5j,n). There was, however, an indication that the magnitude of the semaglutide-induced anorectic effect was attenuated in mice with AP GLP-1R knocked down, compared to that in control mice (Fig. 5q). Also consistent with our earlier studies, semaglutide itself reduced meal size and frequency in mice with intact AP GLP-1R. Reactivation of Sema-TRAP^NTS^ neurons in these mice had a similar effect over the dark phase, particularly on meal size, albeit of lesser magnitude than semaglutide itself (Fig. 5l-m). Conversely, reactivation of Sema-TRAP^NTS^:Dq-*Glp1r*^AP-KD^ neurons had no effect on these satiation or satiety parameters, however, the meal frequency effect of semaglutide itself was fully preserved (Fig. 5o-p).

We then tested whether the negative valence of Sema-TRAP^NTS^ neuron reactivation was also affected by AP GLP-1R knockdown prior to TRAPing. Reactivation in mice with intact AP GLP-1R elicited the same strong avoidance response as semaglutide, as we observed in our previous cohort. However, this avoidance response was largely abolished following reactivation of Sema-TRAP^NTS^:Dq-*Glp1r* ^AP-KD^ neurons (Fig. 5r).

These data reveal that both the satiation and nausea effects of semaglutide, which are recapitulated by reactivation of sema-TRAP^NTS^ neurons, require intact AP GLP-1R. Therefore, they demonstrate that these specific phenotypic components of the drug are mediated by a pathway requiring AP ^GLP-1R^ signalling upstream of the NTS, which is sufficient to acutely elicit bodyweight loss. Conversely, our data demonstrate that semaglutide potentiates satiety both via the same AP^GLP-1R^→NTS pathway recruited for its effects on satiation and nausea, and also via an additional pathway beginning at a site of action outside of the DVC.

### Area postrema GLP-1R are required for semaglutide-induced suppression of motivation to consume Western diet

We next investigated the remaining anorectic phenotype component – suppression of motivation to consume Western diet. We utilised the same cohorts of Sema-TRAP^NTS^:Dq-*Glp1r*^AP-Con^ and Sema-TRAP^NTS^:Dq-*Glp1r*^AP-KD^ mice to test the effect of Sema-TRAP^NTS^ neuron reactivation, and the necessity of AP GLP-1R, on performance in our progressive ratio task (Fig. 6a). In mice with intact AP GLP-1R, reactivation of Sema-TRAP^NTS^ neurons potently reduced the number of nose pokes made, maximum ratio achieved, and number of rewards earned, to a greater extent than semaglutide itself (Fig. 6b-d). Tracking mouse locomotion during this assay revealed a decrease in overall distance travelled and average velocity during the full test session. However, when analysis was restricted to the period in which mice earned an equivalent number of rewards under all three conditions (reward-matched analysis), the suppressive effects of both Sema-TRAP^NTS^ reactivation and semaglutide itself on distance travelled were abolished. This demonstrates that locomotor performance was comparable until the reactivation-induced breakpoint was reached (EDFig. 7a-d). A net loss of bodyweight in this assay was also observed following Sema-TRAP^NTS^ neuron reactivation, albeit to a much more modest extent than that elicited by semaglutide itself in this context (Fig. 6e).

**Figure 7:**
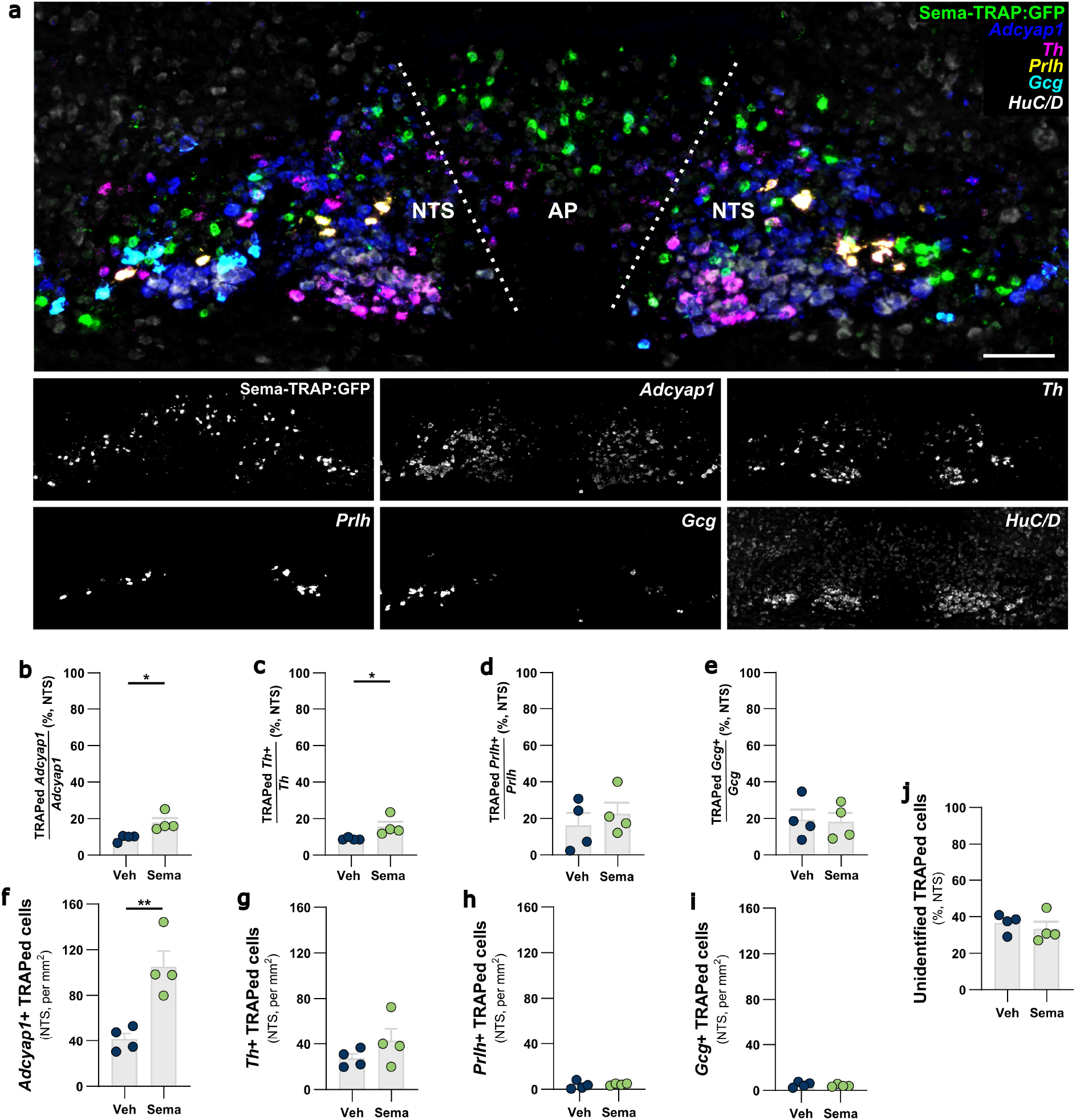
Semaglutide recruits *Adcyap1* and *Th* neurons in the NTS. **a**, representative photomicrograph of coronal DVC section from Sema-TRAP mice showing GFP and HuC/D immunoreactivity and *in situ* hybridisation for *Adcyap1*, *Th*, *Prlh* and *Gcg* mRNA, scale bars = 100μm. **b**, % of *Adcyap1* +ve cells in NTS which were TRAPed: *U*_(4)_=0, *P*=0.0286. **c**, % of *Th* +ve cells in NTS which were TRAPed: *U*_(4)_=0, *P*=0.0286. **d**, % of *Prlh* +ve cells in NTS which were TRAPed: *t*_(6)_=0.706, *P*=0.5067. **e**, % of *Gcg* +ve cells in NTS which were TRAPed: *t*_(6)_=0.1614, *P*=0.8770. **f**, number of *Adcyap1* +ve cells in NTS which were TRAPed: *t*_(6)_=4.31, *P*=0.0051. **g**, number of *Th* +ve cells in NTS which were TRAPed: *t*_(6)_=1.313, *P*=0.2371. **h**, number of *Prlh* +ve cells in NTS which were TRAPed: *t*_(3.4)_=0.4964, *P*=0.6502. **i**, number of *Gcg* +ve cells in NTS which were TRAPed: *t*_(6)_=0.72, *P*=0.5023. **j**, % of TRAPed cells in NTS which were negative for all markers: *t*_(6)_=0.69, *P*=0.5175.

Notably, in mice with AP GLP-1R knocked down before Sema-TRAPing, reactivation of Sema-TRAP ^NTS^ neurons had no effect on any performance parameters in this task, and furthermore the effect of semaglutide itself was abolished (Fig. 6f-h). Similarly, locomotor activity analysis demonstrated an absence of effect of either manipulation on distance and velocity during the full test session, and on reward-matched distance travelled (EDFig. 7e-h). Importantly, while Sema-TRAP^NTS^ neuron reactivation had no effect on bodyweight in this assay, the effect of semaglutide itself on bodyweight was preserved (Fig. 6i), demonstrating the behavioural specificity of the effect of AP GLP-1R knockdown in this context.

We hypothesised that the AP^GLP-1R^-dependent effects on motivation for WD may involve modulation of mesolimbic dopamine signalling, either by inhibition of VTA neurons encoding responses to rewarding stimuli, and/or by activation of neurons responding to aversive stimuli^41,42^. We therefore first quantified immunoreactivity for the neuronal inhibition marker phospho-pyruvate dehydrogenase (pPDH)^43^ in tyrosine hydroxylase (TH) immunoreactive VTA neurons of Sema-TRAP^NTS^:Dq-*Glp1r*^AP-Con^ and Sema-TRAP^NTS^:Dq-*Glp1r*^AP-KD^ mice which were given semaglutide prior to transcardial perfusion (EDFig. 7i). However, pPDH immunoreactivity was only observed in <5% of VTA TH neurons in both groups, there was no effect of AP^GLP-1R^ knockdown, and no correlation between pPDH immunoreactivity and the effect of semaglutide on rewards earned in the progressive ratio task (EDFig. 7j-m). To investigate activation of aversion-responsive neurons, we next quantified immunoreactivity for the neuronal marker of activation, cFos, in VTA neurons of Sema-TRAP^NTS^:Dq-*Glp1r*^AP-Con^ and Sema-TRAP^NTS^:Dq-*Glp1r*^AP-KD^ mice. However, cFos was likewise detected in <5% of VTA TH neurons in both groups, and there was no effect of AP^GLP-1R^ knockdown, and no correlation with the effect of semaglutide on rewards earned in the progressive ratio task (EDFig. 7n-r). Together, these data argue against a change in the somatic activity of VTA dopamine neurons as the basis for the AP^GLP-1R^-dependent reduction in motivation by semaglutide, although modulation of dopamine release in terminal regions cannot be excluded.

### Semaglutide recruits *Adcyap1* and *Th* neurons in the NTS

To identify the molecular identity of semaglutide-recruited NTS neurons, we utilised our sema-TRAPed GFP reporter mice combined with multiplexed *in situ* hybridisation for markers of some major NTS populations previously implicated in eating and sickness behaviours, and response to semaglutide^10,11^, namely *Adcyap1*, *Th*, *Prlh* and *Gcg* (Fig. 7a). Semaglutide increased the proportion of both *Adcyap1*- and *Th*-expressing NTS neurons that were TRAPed, relative to vehicle controls (Fig. 7b,c), whereas the proportions of TRAPed *Prlh* and *Gcg* neurons were unaffected (Fig. 7d,e). This was accompanied by an increase in the overall number of *Adcyap1*+ TRAPed neurons (Fig. 7f), but no significant increase in the number of *Th*+, *Prlh*+, or *Gcg*+ populations (Fig. 7g-i). Semaglutide therefore preferentially recruits *Adcyap1*- and *Th*-expressing NTS neurons, two populations previously implicated in distinct aspects of eating suppression. Notably, approximately one third of all sema-TRAPed neurons in the NTS could not be identified using these markers (Fig. 7j), leaving their identity unresolved.

Overall, by combining our Sema-TRAP^NTS^ reactivation approach with AP-specific knockdown of GLP-1R, we demonstrate that an AP^GLP-1R^→NTS pathway is recruited by semaglutide to potentiate satiation, elicit nausea-like behaviour, and suppress motivation to consume high-fat high-sugar diet. This pathway also mediates part of semaglutide’s ability to potentiate satiety, but the majority of this phenotypic effect appears to be mediated by an alternative pathway starting outside of the DVC. We demonstrate that semaglutide preferentially recruits NTS *Adcyap1*- and *Th*-expressing neurons, broad populations which have previously been implicated in mediating sickness-like responses, non-aversive satiation and/or satiety, and aspects of motivated behaviour. These may therefore contain non-GLP-1R populations which could represent viable targets for tuning the effects profile of next generation GLP-1-based anti-obesity drugs.

## Discussion

Despite the remarkable weight loss efficacy of GLP-1 receptor agonists, including semaglutide, the neuronal populations recruited to mediate their diverse behavioural effects remain poorly resolved. Here, using activity-dependent genetic tagging, population-specific reactivation, and region-specific *Glp1r* knockdown, we identify a common brainstem pathway through which semaglutide coordinates the majority of its behavioural effects. We show that semaglutide engages a substantial population of neurons beyond those expressing GLP-1Rs, particularly within the NTS, and that reactivation of these largely non-GLP-1R NTS neurons specifically recapitulates a subset of the phenotypic components of semaglutide’s anorectic effects. Spatially constrained disruption of *Glp1r* expression in the AP further revealed that these NTS populations are recruited downstream of AP GLP-1 receptors, defining an AP^GLP-1R^→NTS pathway through which semaglutide concurrently potentiates satiation, elicits nausea, and suppresses reward-motivated behaviour.

Previous findings have established the DVC as a major site of action for GLP-1RAs ^5–7,10^, yet the respective contributions of GLP-1R populations in the AP and NTS to specific behavioural components remain incompletely resolved. Both receptor-defined neuron populations are independently capable of reducing food intake and body weight when activated, whereas AP, but not NTS, GLP-1R neurons elicit aversion in conditioned flavour avoidance paradigms^5,7,25^. Their respective contributions to the behavioural effects of semaglutide, however, are less clear. Although ablation of DVC GLP-1R neurons abolishes hypophagia to low-dose semaglutide (10ug/kg)^5^, regional *Glp1r* knockdown or chemogenetic inhibition generally leaves substantial anorectic responses to GLP-1RAs intact^5,7^. AP GLP-1Rs nevertheless appear necessary for the aversive effects of semaglutide^5,7,25^. Our findings reconcile these observations by showing that AP *Glp1r* knockdown does not abolish the overall suppression of food intake by semaglutide, but selectively attenuates semaglutide-induced satiation and nausea, while preserving the majority of its effect on satiety. This aligns with previous evidence that activation of AP GLP-1R neurons suppresses feeding predominantly by promoting satiation and nausea, rather than satiety^5^.

However, whether the AP and NTS GLP-1R neuronal populations represent direct targets which are both substantially activated by semaglutide, and comprise parallel circuits responsible for drug-induced satiation and satiety is unclear, given mixed evidence regarding the full accessibility of NTS GLP-1Rs to peripherally-administered agonists^8,9,32–34,44,45^. By quantifying GLP-1R expression across semaglutide-recruited neurons throughout the rostro-caudal extent of the visceral NTS, we found that direct recruitment of NTS GLP-1R neurons is surprisingly limited, despite extensive recruitment of non-GLP-1R neurons in the NTS. Notably, we performed our sema-TRAP procedure in the dark phase, when circadian regulation of accessibility of the NTS to large blood-borne molecules is most permissive ^46^. The limited direct recruitment we observed is therefore unlikely due to circadian gating of access. Rather, our data suggest that much of the NTS response to semaglutide reflects indirect recruitment, rather than direct engagement of GLP-1R in this region. Consistent with this, reactivation of Sema-TRAP neurons in the NTS primarily recapitulated the satiation and aversive effects of semaglutide, alongside a partial effect on satiety, rather than the full satiety response previously ascribed to the NTS GLP-1R population^5^. This satiety effect is unlikely to arise from the small proportion of NTS GLP-1R neurons captured by Sema-TRAP, since GLP-1R neurons were not significantly recruited over vehicle-TRAP, and reactivation no longer elicited satiety in AP *Glp1r* knockdown mice despite NTS GLP-1Rs being left intact. Furthermore, the satiation and aversion elicited from reactivation, and by semaglutide itself, were effectively abolished by knockdown of AP *Glp1r* expression, whereas a substantial satiety effect of semaglutide persisted. Together these findings implicate a predominantly non-GLP1R NTS population downstream of AP GLP-1Rs as the primary DVC pathway recruited by semaglutide, confirming and extending recent findings^6,7,10^. They further establish that semaglutide drives satiety through at least two distinct sites: this AP^GLP-1R^→NTS pathway, and a second site outside the DVC that remains intact following AP *Glp1r* knockdown. The reduced magnitude of semaglutide’s satiety effect in knockdown animals is consistent with these sites acting in combination.

Beyond its effects on homeostatic eating behaviours, semaglutide suppressed reward-motivated behaviour, consistent with emerging evidence for effects on both food and non-food rewards ^3,4,15^. Although GLP-1R populations within the mesolimbic and forebrain regions have been implicated in reward processing^3,16–18^, whether these populations are directly engaged by semaglutide remains unclear. Our findings instead identify the AP^GLP-1R^→NTS pathway as a crucial upstream mediator of semaglutide-induced suppression of eating motivation. Reactivation of Sema-TRAP^NTS^ neurons was sufficient to suppress reward-motivated behaviour, while AP *Glp1r* knockdown abolished this effect of semaglutide. Previous studies similarly place several midbrain and forebrain regions downstream of semaglutide-responsive DVC populations, including the parabrachial nucleus (PBN) and central amygdala (CeA)^5,10^, providing potential routes through which semaglutide’s action in the DVC could influence motivated behaviour.

An important unresolved question is whether reward suppression and aversion represent parallel outputs from the AP GLP-1R site of action, or whether they are mechanistically coupled. Given that AP GLP-1R signalling was required for both effects, reduced motivation for palatable food could arise, at least in part, from the negative valence state induced by semaglutide. Alternatively, AP GLP-1R-dependent recruitment of distinct NTS populations may engage parallel downstream pathways that independently mediate aversion and motivation. Our molecular characterisation of sema-TRAP^NTS^ neurons identifies candidate populations for such a dissociation. Semaglutide recruited NTS *Adcyap1* neurons, consistent with recent work implicating this population and its hypothalamic projections in the drug’s effects on food intake and body weight^10,47^, and with their established role in sickness-related anorexia^11,48^. Semaglutide also recruited NTS *Th* neurons, which suppress cue-induced appetitive behaviour and cue-evoked dopamine signalling without inducing malaise^49^, whereas *Prlh* and *Gcg* populations were not recruited above vehicle. NTS *Adcyap1*- and *Th*-expressing populations therefore offer plausible candidates for the potentially dissociable aversive and reward-suppressing components of semaglutide’s actions. Both are themselves heterogeneous^47,50^, however, and a substantial proportion of Sema-TRAP^NTS^ neurons did not express any of the markers examined. Defining whether these populations independently mediate aversion and reward suppression will be crucial for establishing whether the motivation-modifying benefits of GLP-1RAs can be dissociated from their adverse effects.

There are some inherent limitations to our study, including that we cannot definitively determine to what extent the NTS neurons stochastically active under our TRAPing conditions when semaglutide was not administered (Veh-TRAP^NTS^) overlap with those recruited as part of the mechanism of action of semaglutide. It is certainly plausible that there is at least some degree of overlap between these populations, given the qualitative similarities between the behavioural and metabolic phenotypes caused by their reactivation, including transient suppression of food intake and a modest increase in satiety. Nevertheless, several observations support the specificity of our Sema-TRAP^NTS^ model and the behavioural phenotype caused by Sema-TRAP^NTS^ neuron reactivation. A recent study using a similar semaglutide-induced TRAPing approach reported comparable effects following reactivation of semaglutide-recruited neurons in the NTS and AP combined, but a lack of detectable phenotype following reactivation of vehicle-TRAPed neurons^10^. Our approaches were quite similar, including the dose of semaglutide used as the TRAPing stimulus, but subtle methodological differences could potentially account for the modest Veh-TRAP^NTS^ effects observed in our study. Most compellingly, our AP *Glp1r* knockdown effectively abolished the anorectic effects of subsequent Sema-TRAP^NTS^ reactivation. This behavioural specificity clearly demonstrates that the Sema-TRAP^NTS^ population is recruited downstream of AP GLP-1Rs. Any residual effects observed in individual animals may reflect variability in the stochastic activity-dependent neuronal tagging, and/or the extent of *Glp1r* knockdown achieved between animals.

Together, our findings identify an AP^GLP-1R^→NTS circuit as a key pathway through which semaglutide coordinates satiation, nausea and reward-motivated behaviour, and highlight the importance of neuronal populations recruited downstream of the receptor itself in shaping GLP-1RA actions. We show that these predominantly non-GLP-1R NTS neurons include *Adcyap1*- and *Th*-expressing populations with distinct reported roles in malaise and appetitive behaviour. Resolving whether they constitute parallel aversive and non-aversive pathways, and how they relate to vagal afferent signalling, will be important next steps. Moreover, the persistence of semaglutide effects following AP *Glp1r* knockdown points to additional sites and mechanisms of action that remain to be defined. Resolving this distributed circuitry may ultimately provide opportunities to dissociate the therapeutically beneficial effects of GLP-1RAs from adverse effects such as nausea.

## Methods

### Animals

Adult mice (8-35 weeks of age; both sexes) were maintained on a 12hr light/dark cycle at 22-24 °C and 45-65% relative humidity, with *ad libitum* access to standard rodent chow (Teklad 2018) and tap water. Mice were group-housed until surgery and/or behavioural experiments. All animals were on a C57BL/6J genetic background, and included C57BL/6J wild type mice (Charles River), FosTRAP/TRAP2 mice (*Fos^tm^*^2^*^.1(icre/ERT2)Luo^*/J; JAX 030323), FosTRAP x Ai9 mice (*Fos^tm2.1(icre/ERT2)Luo^*/J x Ai9; JAX 030323 x JAX 007909), and FosTRAP x NuTRAP mice (*Fos^tm2.1(icre/ERT2)Luo^*/J x B6;129S6-*Gt(ROSA)26Sor^tm2(CAG-NuTRAP)Evdr^*/J; JAX 030323 x JAX 029899). Before experiments, mice were habituated for 3-5 days to handling, injections, and behaviour chambers, as appropriate. Animals were randomly assigned to experimental groups where appropriate, and balanced for sex and age as far as possible. No statistical methods were used to predetermine sample size. Sample sizes were based on previous studies and laboratory experience.

All experiments were conducted in the UK in accordance with the U.K. Animals (Scientific Procedures) Act 1986, and experimental protocols were approved by the UCL Animal Welfare and Ethical Review Body (Bloomsbury Campus).

### TRAP Induction

FosTRAP mice were fasted for 5hr before TRAP stimuli. At dark onset, mice received an intraperitoneal (i.p.) injection of either vehicle or semaglutide (see ***Drugs***) in their home cage. Three hour later, mice received an i.p. injection of 4-hydroxytamoxifen (4-OHT; 30 mg/kg; 3 mL/kg; Sigma, H6278) dissolved in Chen oil (1:4 castor oil (Sigma, 259853): sunflower seed oil (Sigma, S5007)) ^51^. Standard chow was returned 3hr after 4-OHT administration.

### Drugs

Semaglutide (Cayman Chemical, 29969) was dissolved in 0.2% DMSO and diluted in sterile phosphate-buffered saline (PBS) immediately before use. Vehicle consisted of 0.2% of DMSO in PBS. Deschloroclozapine (DCZ; Hello Bio, HB9126) was dissolved in sterile saline. Unless otherwise stated, semaglutide was administered at 60 µg/kg (5 mL/kg, i.p.), vehicle at 5 mL/kg (i.p.), and DCZ at 0.1 mg/kg (2 mL/kg, i.p.).

### Stereotaxic Surgery

FosTRAP mice were anaesthetized with ketamine hydrochloride (100 mg/kg; Vetalar) + medetomidine (1 mg/kg; Dormitor) i.p., and given meloxicam analgesia (5mg/kg) subcutaneous (s.c.) injection. Appropriate depth of surgical anaesthesia was determined by absence of pedal reflex, and animal temperature was maintained using a homeothermic monitoring system. Mice were placed in a stereotaxic frame and the head flexed downwards such that the nose and neck were at a right angle. The scalp was incised from the occipital crest to first vertebrae, and muscle layers parted to expose the atlanto-occipital membrane. The membrane was bisected horizontally with a 1 mm blade Phaco knife (PFM, 1001048) to expose the dorsal brainstem. Viral vectors were administered via pulled glass micropipettes into the AP (+0.65 mm AP, 0 mm ML, and -0.1 mm DV) or bilaterally into the NTS (+0.2 mm AP, ±0.27 mm ML, and -0.2 DV) from obex.

For chemogenetic activation studies, 25 nL of AAV9-hSyn-DIO-hM3Dq-mCherry (1×10^12^; Addgene, 44361-AAV9) or as a control AAV5-hSyn-DIO-mCherry (1×10^12^; Addgene, 50459-AAV5) were bilaterally injected into the NTS. To knockdown *Glp1r*, 25 nL of custom produced AAV8/2-hSyn-shGL1R-mScarlet (8×10^11^) or as a control AAV9-hSyn-[4x:sh(NS)]-EGFP (8×10^11^; ETHZ, v308-9) was injected into the AP.

Mice were treated daily with meloxicam (5mg/kg, s.c.) for 48 hours for post-operative analgesia, and recovered for ≥2 wks (≥3 wks for *Glp1r* knockdown) before TRAP induction. Behavioural testing or tissue collection was performed ≥2 wks after TRAP induction.

### Metabolic Phenotyping

#### Experimental Procedures

Metabolic phenotyping was performed using the Promethion metabolic phenotyping system (Sable Systems International), allowing continuous measurement of feeding behaviour, indirect calorimetry and locomotor activity. Mice were habituated to the system before experimentation and maintained under the same environmental conditions described above, with food access automatically restricted for the last two hours of light phase daily, to entrain consistent dark onset feeding without inducing negative energy balance. Semaglutide, DCZ or vehicle was administered 30 min before the onset of the dark phase, and metabolic parameters were recorded continuously for 22hr. The order of drug conditions were counterbalanced and separated by a 72hr washout period where mice had *ad libitum* access to food.

#### Data Processing & Analysis

All raw data from Promethion system were initially processed using MacroInterpreter software v25.2, running OneClickMacro v2.53.3 (Sable Systems Inc). These datasets were further processed using custom Python scripts to extract the data from the relevant 22hr time windows commencing with dark onset food access immediately after drug dosing, reordered by drug treatment. Metabolic phenotyping data were analysed and presented in accordance with the consensus guide for indirect calorimetry^52^, adapted where necessary for relevance to the within-subjects crossover experimental designs used. Meal pattern analysis parameters were based on our previous empirical optimisation of these for mouse feeding behaviour^53^.To minimise overestimation of food intake due to caching and/or false positive sensor readings, food intake bouts with an intake rate >0.1 g/min were removed prior to analysis of hourly intakes and meal pattern parameters. Non-cumulative hourly food intake data were analysed to identify the time window during which drugs actively affected intake. This window was then used for subsequent analysis of meal patterns and other relevant parameters including energy expenditure. Given the potential for sustained effects on metabolic phenotyping parameters beyond the acute active drug effect windows, key parameters were additionally analysed across the dark and light phases of the circadian cycle. Energy expenditure was analysed with bodyweight on each day of dosing as a covariate using custom Python scripts. Where the treatment x BW interaction was non-significant, data were analysed by ANCOVA, otherwise by GLM. Separate to these analyses, energy expenditure, respiratory exchange ratio, and locomotor activity data were also extracted in 5 minute time bins and plotted as 30 minute rolling averages to aid visualisation over the full circadian cycle.

### Western Diet Intake

Mice were habituated to a custom-built HM2++ feeding system, based on the HM2 system (MBRose)^23^, housed within standard commercial operant chambers (Med Associates). During habituation and experimental sessions, mice were fasted for the final 5hr of the light phase before receiving *ad libitum* access to western diet (Research Diets, D12079B) for 4hr from dark onset. On experimental days, semaglutide or vehicle was given 2hr prior to food access. Food intake and meal pattern data was measured by HM02 software (MBRose), and the difference in manually weighed loaded hopper before and after the feeding session used to validate total intake output in case of system error. Feeding bouts are defined by the system as intake > 0.02 g, and applies an inter-bout interval of 30 sec. Meal pattern analysis was processed from bouts, and defined as > 0.04 g with an inter-meal interval > 180 sec.

### Conditioned Flavour Avoidance

Conditioned flavour avoidance was used to assess avoidance of a novel tastant following drug pairing as an index of aversive responses. Mice were water restricted for ∼16hr and habituated to a two-bottle choice paradigm. On pairing day, mice consumed 0.15% saccharin + 0.05% cherry Kool-Aid (Fig. 1) or 5% sucrose (Fig. 4+5) followed by a DCZ or semaglutide injection. On test day, mice were presented with saccharin or sucrose and water for 1hr. A preference ratio was calculated using the equation: Preference ratio = mass of sucrose solution consumed / total mass of liquids consumed.

### Progressive Ratio Operant Task

Operant testing was conducted during the light phase in the HM2++ containing operant chambers described above, with the house light illuminated. Mice were fasted for 18hr before each behavioural session. Rewards consisted of western diet and were delivered from the HM2++ hopper, located behind a guillotine door on the wall opposite the nose-poke port; intake was recorded by the HM02 software. The task comprised three phases: reward learning, progressive ratio training, and pharmacological testing. During reward learning, food was available for 20s every 30s independent of nose poke responses, and reward availability was signalled by a 5 kHz tone. Mice were then trained on a fixed-ratio (FR1) schedule, in which each rewarded nose poke was accompanied by a 2 kHz auditory cue. Reward access was initially 12s and was reduced to 7s as performance improved ^54^. Once mice earned 60 rewards within a session^54^, they received saline injections (5 ml/kg, i.p.) 60 min before testing and were advanced to an FR5 schedule. Mice consumed an average of ∼20 mg WD per reward earned, equivalent to the mass of pellets used in standard operant systems (EDFig. 2d).

Under the PR schedule, the response requirement increased exponentially according to the schedule *F(i)* = 5e^0.2i^–5, where *F(i)* represents the number of nose pokes required to obtain reward *i*^55^. Sessions were 60min long, unless terminated earlier due to 10min without a nose poke^56^, and the highest completed ratio (breakpoint) was taken as a measure of reward motivation. Before testing, mice completed at least two PR training sessions and were required to demonstrate a stable baseline, defined as a breakpoint varying by less than 20% between the final two sessions.

Latency to first nose poke was measured from session onset. Post-reinforcement pause (PRP) was defined as the interval from the end of reward access to the first nose poke of the next ratio. Working time was defined as the interval from the first to the last nose poke of each ratio. For each mouse, PRP and working time were analysed only for rewards obtained under both vehicle and semaglutide treatment.

On test days, mice were given vehicle or semaglutide 2hr before, or DCZ 15min before behavioural testing. Treatment order was counterbalanced, and test sessions were separated by a 72hr washout period during which mice had *ad libitum* access to food.

### Video Analysis

#### Markerless pose estimation and movement analysis

Complete PR sessions were recorded on overhead USB 3.1 monochrome cameras (Imaging Source 37AUX273) at 30 frames per second (1,440 x 1,080 pixels). Pose estimation was performed using SLEAP (Social LEAP Estimated Animal Pose, v1.5.1) framework^57^. A single-animal model was trained to track four keypoints: nose, left ear, right ear and tail base. To ensure generalisation across chambers, the model was trained on videos of habituation sessions recorded in eight operant chambers using an iterative human-in-the-loop approach. An initial model was trained on 500 manually labelled frames from one chamber and used to predict a further 500 frames, which were corrected and added to the training set. For each subsequent chamber, the current model was first evaluated on 500 frames before annotation; these frames were then corrected and used for retraining, and performance was reassessed on a second set of 500 frames. The final model was trained on 8,000 labelled frames (1,000 per chamber). The model was trained using a single instance U-Net model with max stride 16, 16 filters and ± 180° rotation. Model performance was evaluated against manually annotated ground truth using the instance score, keypoint detection precision, recall and F1 score, and localisation error (Euclidean distance in pixels between predicted and annotated keypoints). The model achieved a keypoint detection precision score of 0.997, recall score of 0.996, an F1 score of 0.996 and a localisation error of 7.06px/1.96mm. Pose data were analysed using SimBA (Simple Behavioural Analysis^58^ v.4.9.9). Keypoint trajectories were smoothed using Savitzky-Golay method, and pixel coordinates were converted to centimetres using the known chamber dimensions of 18.2cm. Distance travelled and velocity were calculated from the nose node, and path plots were generated to visualise spatial behaviour.

#### Behavioural syllable analysis

Behavioural syllables were identified using Keypoint-MoSeq (v.0.6.6) applied to all four tracked keypoints (nose, left ear, right ear, tail base) from 11 mice across both treatment sessions. Keypoints with a SLEAP confidence score below 0.5, or located more than five times the median distance from the body centre, were interpolated and keypoint error priors were calibrated using 20 manually annotated frames. Poses were aligned along the nose-tail axis, after which four latent dimensions explained 90% of pose variance. To determine the appropriate timescale for behavioural segmentation, we performed a κ scan across all tracking data and selected the value that yielded a median syllable duration of approx. 400 ms, consistent with previous mouse studies^59^. The final model (κ = 5 x 10^9^) identified 90 syllables, of which 21 each comprised at least 0.5% of frames were retained for analysis. Analysis was performed as reward-matched analysis, both sessions for each mouse were truncated at the final reward of the session with fewer rewards (typically semaglutide).

To assess overall shifts in behavioural composition, principal component analysis (PCA) was performed on the proportion of frames spent in each retained syllable for each mouse and session, with each syllable standardised to zero mean and unit variance. The sign of PC1 was orientated so that semaglutide-induced shifts were in a consistent direction.

Retained syllables were manually annotated by two raters. For each syllable, raters viewed a grid of 64 randomly sampled instances drawn across mice and treatments so raters were blind to condition. A behavioural label was assigned when more than 70% of instances showed concordant behaviour. Seventeen labelled syllables were then grouped into 6 behavioural categories. The proportion of time in each category was calculated as the category’s frames divided by all labelled frames in the window (%), with unclassified frames shown as “other”. Total time was calculated as frames divided by 30 fps.

### Immunofluorescence and In-situ Hybridization

#### Tissue Preparation

In preparation for immunofluorescent staining, mice were deeply anaesthetised then transcardially perfused with ice-cold heparinised PBS then 4% paraformaldehyde (PFA) in PBS. Brains and nodose ganglia (when required) were extracted and post-fixed in 4% PFA at 4 °C overnight (≤2hr for nodose ganglia), before being cryoprotected in 30% sucrose solution for ≥24 hr at 4 °C. Brains were sectioned into 25 μm coronal sections, collected free-floating and stored at 4 °C until processing for immunofluorescence labelling. Nodose ganglia were sectioned into 10 μm sections, collected directly onto Superfrost Plus microscope slides and stored at −20 °C until processing for immunofluorescence labelling. For *in situ* hybridisation, mice were deeply anaesthetised then decapitated. Brains were quickly removed, immediately frozen in optimal cutting temperature compound (OCT) on dry ice and stored at -80 °C until processing by RNAscope.

#### Immunofluorescence

Brain sections were processed for immunofluorescent detection of fluorescent reporters (eYFP, eGFP and RFP), cell-type marker Tyrosine hydroxylase, the neuronal marker HuC/D, the vagal afferent marker PHOX2B, and the markers of neuronal activity and inhibition cFOS and pPDH, using previously validated protocols (Brierley, 2021). For all immunofluorescent studies, except those performed in conjunction with RNAscope, sections were blocked in PBS containing 2.5% normal goat and/or donkey serum and 0.1% Triton X-100 (1hr, room temperature), followed by incubation with primary antibodies (>16 hr, 25° C), and fluorophore-conjugated secondary antibodies (2 hr, room temperature). cFOS immunolabelling was instead amplified using tryamide signal amplification (TSA), with an HRP-conjugated anti-rabbit secondary antibody (2hr, room temperature), followed by TSA Vivid 650 (5min). Sections were mounted onto SuperFrost Plus slides with CitiFluor AF1 mounting medium and imaged.

#### RNAscope In Situ Hybridisation

*In situ* hybridisation was performed using the RNAscope Multiplex Fluorescent Kit v2 according to the manufacturer’s instructions. Sections were hybridised with probes targeting *Glp1r* (Bio-techne, 418851-C2), *Th* (info), *Gcg* (info), *Prlh* (info) and *Adcyap1* (info), detected using Opal dyes (Quanterix), followed by immunofluorescent detection of HuC/D and GFP as described above.

#### Image Acquisition and Analysis

For TRAP quantification and cFOS or pPDH analysis, brain sections were imaged using the EVOS M7000 Imaging System (Thermo Fisher). Widefield, z-stacks were acquired at 20× magnification in DAPI, GFP, RFP, and Cy5 channels and converted to 2D maximum-intensity projections using FIJI.

For 2D RNAscope analysis, images were acquired using an Akoya PhenoImager HT 2.0 (Quantrix) at 20x magnification. Sections hybridised with negative-control probes were used to determine spectral unmixing parameters. No post-acquisition modifications were applied during quantification. *Glp1r, Th, Gcg, Prlh and Adcyap1* RNAscope quantification was performed using QuPath (version 0.7.0) and custom scripts. For quantification of *Glp1r* knockdown, background regions of interest were measured and their mean fluorescence intensity subtracted from *Glp1r* signal intensity.

For 3D RNAscope analysis, confocal z-stacks were acquired using a Nikon NSPARC confocal microscope at 25× silicon-oil immersion objective. Sections hybridised with negative-control probes were used to determine acquisition parameters, and channels were acquired sequentially. Z-stacks were acquired at 5μm intervals, with no post-acquisition modifications applied before quantification. 3D quantification was performed using Imaris (Oxford Instruments, version 11), using the Surface, Spots and Cell functions. Batch analysis parameters were applied consistently across images to identify HuC/D+ neurons and GFP+ TRAPed neurons, and to quantify *Glp1r* RNAscope puncta within individual cells.

### Statistical analysis and data presentation

All data are presented as mean ± standard error of the mean. Results were considered statistically significant when α < 0.05. All significant pairwise comparisons or ANOVA main effects are shown on all plots, near-significant pairwise comparisons (*P* = 0.05 – 0.1) are shown with exact P values, with all non-significant pairwise comparisons (*P* ≥ 0.1) omitted from plots for visual clarity. Results of all statistical tests performed are given in accompanying figure legends. Sample sizes (split by sex) are given in experimental design schematic figures for each study. Where data from individual animals were omitted from specific analyses (e.g. due to unavailability of data due to technical errors), this is reflected in lower degrees of freedom values in statistical test details in figure legends.

Statistical testing was performed using Graphpad Prism v11.0.1 or custom scripts in Python. Comparisons between 2 conditions were performed using paired or unpaired Student’s t-tests, or non-parametric equivalents where appropriate, and are all two-tailed unless otherwise specified in figure legends. Comparisons between 3 or more unpaired conditions were performed by one-way ANOVA, or for paired data by one-way repeated measures ANOVA (with Greenhouse-Geiser correction where sphericity was violated, indicated by epsilon <0.75), or mixed-effects model (REML) if there were missing values. Comparisons between 2 or more conditions for 2 variables were performed by two-way ANOVA, two-way repeated measures ANOVA, mixed-model ANOVA, or mixed-effects model (REML) as appropriate. Where significant ANOVA main effects or interactions were observed, post-hoc pairwise comparisons were made with Holm-Sidak corrections for multiple comparisons.

## Acknowledgements

We would like to thank Sabrina Goldring, Jennifer Lee, Bidisha Roy, Jonathan Aderanti, Shirley Nannono-Senabulya and Sophie McManus for their invaluable contributions conducting preliminary method optimisation and validation studies for several approaches used in this study. We additionally thank Virginia Silio and Alan Grieg (UCL Biosciences imaging facility) and Jiten Manji (UCL Cancer Institute) for invaluable support with microscopy.

## Funding

This work was funded by a Wellcome Trust / Royal Society Sir Henry Dale Fellowship (223279/Z/21/Z) and British Society of Neuroendocrinology project support grant to DIB. ST was funded by the MRC (MR/X003604/1).

## Declaration of Interests

A.E.A. is now an employee and shareholder at AstraZeneca. The other authors declare no conflicts of interest.

## Notes

### Summary of Updates

Additional data have been added and figures and text updated throughout.

## References

1. Drucker, D. J. Efficacy and Safety of GLP-1 Medicines for Type 2 Diabetes and Obesity. Diabetes Care dci240003 (2024) doi:10.2337/dci24-0003.

2. Jones, L. A. & Brierley, D. I. GLP-1 and the Neurobiology of Eating Control: Recent Advances. Endocrinology 166, bqae167 (2025).

3. Godschall, E. N. et al. A brain reward circuit inhibited by next-generation weight-loss drugs in mice. Nature 1–10 (2026) doi:10.1038/s41586-026-10444-4.

4. Drucker, D. J. The benefits of GLP-1 drugs beyond obesity. Science 385, 258–260 (2024).

5. Huang, K.-P. et al. Dissociable hindbrain GLP1R circuits for satiety and aversion. Nature 1–9 (2024) doi:10.1038/s41586-024-07685-6.

6. Gao, C. et al. Semaglutide drives weight loss through cAMP-dependent mechanisms in GLP1R-expressing hindbrain neurons. Nat Metab 8, 1330–1349 (2026).

7. Yacawych, W. T. et al. A single dorsal vagal complex circuit mediates the aversive and anorectic responses to GLP1R agonists. https://www.jci.org/articles/view/193743/pdf (2026) doi:10.1172/JCI193743.

8. Adams, J. M. et al. Liraglutide modulates appetite and body weight through glucagon-like peptide 1 receptor-expressing glutamatergic neurons. Diabetes 67, 1538–1548 (2018).

9. Costa, A. et al. Anorectic and aversive effects of GLP-1 receptor agonism are mediated by brainstem cholecystokinin neurons, and modulated by GIP receptor activation. Molecular Metabolism 55, 101407 (2022).

10. Teixidor-Deulofeu, J. et al. Semaglutide effects on energy balance are mediated by Adcyap1+ neurons in the dorsal vagal complex. Cell Metabolism 0, (2025).

11. Wang, H. et al. Parallel gut-to-brain pathways orchestrate feeding behaviors. Nat Neurosci 1–16 (2024) doi:10.1038/s41593-024-01828-8.

12. Dowsett, G. K. C. et al. A survey of the mouse hindbrain in the fed and fasted states using single-nucleus RNA sequencing. Molecular Metabolism 53, 101240 (2021).

13. Ludwig, M. Q. et al. A cross-species atlas of the dorsal vagal complex reveals neural mediators of the effects of cagrilintide on energy balance. Nat Metab 8, 1350–1367 (2026).

14. Ludwig, M. Q., Todorov, P. V., Egerod, K. L., Olson, D. P. & Pers, T. H. Single-Cell Mapping of GLP-1 and GIP Receptor Expression in the Dorsal Vagal Complex. Diabetes 70, 1945–1955 (2021).

15. Blundell, J. et al. Effects of once-weekly semaglutide on appetite, energy intake, control of eating, food preference and body weight in subjects with obesity. *Diabetes*, Obesity and Metabolism 19, 1242–1251 (2017).

16. Merkel, R. et al. An endogenous GLP-1 circuit engages VTA GABA neurons to regulate mesolimbic dopamine neurons and attenuate cocaine seeking. Science Advances 11, eadr5051 (2025).

17. Edvardsson, C. E., Cadeddu, D., Ericson, M., Adermark, L. & Jerlhag, E. An inhibitory GLP-1 circuit in the lateral septum modulates reward processing and alcohol intake in rodents. eBioMedicine 115, (2025).

18. Duran, M., Zeng, N., Cutts, E. J., Habegger, K. & Hardaway, J. A. The central amygdala integrates exogenous glucagon-like peptide 1 signals. 2026.04.06.716705 Preprint at 10.64898/2026.04.06.716705 (2026).

19. Kim, K. S. et al. GLP-1 increases preingestive satiation via hypothalamic circuits in mice and humans. Science 385, 438–446 (2024).

20. Brierley, D. I. et al. Central and peripheral GLP-1 systems independently suppress eating. Nature Metabolism 2021 3, 258–273 (2021).

21. Shah, H. & Ayala, J. E. Prolonged Semaglutide Treatment Reveals Stage-Dependent Changes to Feeding Behavior and Metabolic Adaptations in Male Mice. Diabetes 75, 288–300 (2025).

22. Börchers, S. & Skibicka, K. P. GLP-1 and Its Analogs: Does Sex Matter? Endocrinology 166, bqae165 (2025).

23. Rathod, Y. D. & Fulvio, M. D. The feeding microstructure of male and female mice. PLOS ONE 16, e0246569 (2021).

24. Borner, T. et al. GIP Receptor Agonism Attenuates GLP-1 Receptor Agonist-Induced Nausea and Emesis in Preclinical Models. Diabetes 70, 2545–2553 (2021).

25. Zhang, C. et al. Area Postrema Cell Types that Mediate Nausea-Associated Behaviors. Neuron 109, 461–472.E5 (2020).

26. Drucker, D. J. The benefits of GLP-1 drugs beyond obesity. Science 385, 258–260 (2024).

27. Wang, W. et al. Associations of semaglutide with incidence and recurrence of alcohol use disorder in real-world population. Nat Commun 15, 4548 (2024).

28. Hodos, W. Progressive ratio as a measure of reward strength. Science 134, 943–944 (1961).

29. Sharma, S., Hryhorczuk, C. & Fulton, S. Progressive-ratio responding for palatable high-fat and high-sugar food in mice. J Vis Exp e3754 (2012) doi:10.3791/3754.

30. Alvarez-Sekely, C. S. et al. Comparison of progressive hold and progressive response schedules of reinforcement. Behavioural Processes 205, 104822 (2023).

31. Bradshaw, C. M. & Killeen, P. R. A theory of behaviour on progressive ratio schedules, with applications in behavioural pharmacology. Psychopharmacology 222, 549–564 (2012).

32. Ast, J. et al. Super-resolution microscopy compatible fluorescent probes reveal endogenous glucagon-like peptide-1 receptor distribution and dynamics. Nature Communications 11, 1–18 (2020).

33. Gabery, S., et al. Semaglutide lowers body weight in rodents via distributed neural pathways. JCI Insight 5, e133429 (2020).

34. Secher, A. et al. The arcuate nucleus mediates GLP-1 receptor agonist liraglutide-dependent weight loss. The Journal of clinical investigation 124, 4473–88 (2014).

35. Allen, W. E. et al. Thirst-associated preoptic neurons encode an aversive motivational drive. Science 357, 1149–1155 (2017).

36. DeNardo, L. A. et al. Temporal evolution of cortical ensembles promoting remote memory retrieval. Nature Neuroscience 22, 460–469 (2019).

37. Ambler, M., Hitrec, T., Wilson, A., Cerri, M. & Pickering, A. Neurons in the Dorsomedial Hypothalamus Promote, Prolong, and Deepen Torpor in the Mouse. J. Neurosci. 42, 4267–4277 (2022).

38. McDougle, M. et al. Separate gut-brain circuits for fat and sugar reinforcement combine to promote overeating. Cell Metabolism 36, 393–407.e7 (2024).

39. Roh, H. C. et al. Simultaneous Transcriptional and Epigenomic Profiling from Specific Cell Types within Heterogeneous Tissues In Vivo. Cell Reports 18, 1048–1061 (2017).

40. Kupari, J., Häring, M., Agirre, E., Castelo-Branco, G. & Ernfors, P. An Atlas of Vagal Sensory Neurons and Their Molecular Specialization. Cell Reports 27, 2508–2523.e4 (2019).

41. Prévost, E. D. et al. Distinct properties of glutamate-dopamine, nonglutamate-dopamine, and glutamate-only ventral tegmental area neurons. Science Signaling 19, eaee2028 (2026).

42. Brischoux, F., Chakraborty, S., Brierley, D. I. & Ungless, M. A. Phasic excitation of dopamine neurons in ventral VTA by noxious stimuli. Proceedings of the National Academy of Sciences of the United States of America 106, 4894–9 (2009).

43. Yang, D. et al. Phosphorylation of pyruvate dehydrogenase inversely associates with neuronal activity. Neuron 112, 959–971.e8 (2024).

44. Trapp, S. & Brierley, D. I. Brain GLP-1 and the regulation of food intake: GLP-1 action in the brain and its implications for GLP-1 receptor agonists in obesity treatment. British Journal of Pharmacology 179, 557–570 (2022).

45. Ørskov, C., Poulsen, S. S., Mørten, M. & Holst, J. J. Glucagon-Like Peptide I Receptors in the Subfornical Organ and the Area Postrema Are Accessible to Circulating Glucagon-Like Peptide I. Diabetes 45, 832–835 (1996).

46. Chrobok, L. et al. Timekeeping in the hindbrain: a multi-oscillatory circadian centre in the mouse dorsal vagal complex. Commun Biol 3, 1–12 (2020).

47. Sköldheden, S. B., Teixidor-Deulofeu, J., Ruud, J. & Ruud, L. E. Semaglutide engages distinct brainstem-to-hypothalamus circuits to suppress motivated feeding and regulate ketogenesis and energy expenditure. 2026.09.04.749561 Preprint at 10.64898/2026.09.04.749561 (2026).

48. Ilanges, A. et al. Brainstem ADCYAP1+ neurons control multiple aspects of sickness behaviour. Nature 1–11 (2022) doi:10.1038/s41586-022-05161-7.

49. Yap, J. A., Gu, Y. H., Vuong, C., Power, J. M. & Ong, Z. Y. Nucleus of the solitary tract regulation of cue-induced appetitive behaviors via midbrain dopamine neurons. Neuropsychopharmacol. 1– 10 (2026) doi:10.1038/s41386-026-02417-y.

50. Hes, C. et al. A unified rodent atlas reveals the cellular complexity and evolutionary divergence of the dorsal vagal complex. eLife 14, (2025).

51. Guenthner, C. J., Miyamichi, K., Yang, H. H., Heller, H. C. & Luo, L. Permanent genetic access to transiently active neurons via TRAP: Targeted recombination in active populations. Neuron 78, 773–784 (2013).

52. Banks, A. S. et al. A consensus guide to preclinical indirect calorimetry experiments. Nat Metab 7, 1765–1780 (2025).

53. Jiang, W. et al. Brainstem GLP-1 neurons modulate physiological satiation and drive sustained weight loss in obese mice. Molecular Metabolism 107, 102347 (2026).

54. Cambre, E., Christenfeld, E., Torres, A. H. & Canetta, S. Measuring Motivation Using the Progressive Ratio Task in Adolescent Mice. Current Protocols 3, e776 (2023).

55. Ghidewon, M. et al. Growth differentiation factor 15 (GDF15) and semaglutide inhibit food intake and body weight through largely distinct, additive mechanisms. *Diabetes*, Obesity and Metabolism 24, 1010–1020 (2022).

56. Johnson, A. R., Christensen, B. A., Kelly, S. J. & Calipari, E. S. The influence of reinforcement schedule on experience-dependent changes in motivation. J Exp Anal Behav 117, 320–330 (2022).

57. Pereira, T. D. et al. SLEAP: A deep learning system for multi-animal pose tracking. Nat Methods 19, 486–495 (2022).

58. Goodwin, N. L. et al. Simple Behavioral Analysis (SimBA) as a platform for explainable machine learning in behavioral neuroscience. Nat Neurosci 27, 1411–1424 (2024).

59. Weinreb, C. et al. Keypoint-MoSeq: parsing behavior by linking point tracking to pose dynamics. Nat Methods 21, 1329–1339 (2024).

